# Viral-mediated ubiquitination impacts interactions of host proteins with viral RNA and promotes viral RNA processing

**DOI:** 10.1101/2020.06.05.136671

**Authors:** Christin Herrmann, Joseph M. Dybas, Jennifer C. Liddle, Alexander M Price, Katharina E. Hayer, Richard Lauman, Caitlin E. Purman, Matthew Charman, Eui Tae Kim, Benjamin A Garcia, Matthew D Weitzman

## Abstract

Viruses promote infection by hijacking host ubiquitin machinery to counteract or redirect cellular processes. Adenovirus encodes two early proteins, E1B55K and E4orf6, that together co-opt a cellular ubiquitin ligase complex to overcome host defenses and promote virus production. Adenovirus mutants lacking E1B55K or E4orf6 display defects in viral RNA processing and protein production, but previously identified substrates of the redirected ligase do not explain these phenotypes. Here we used a quantitative proteomics approach to identify substrates of E1B55K/E4orf6-mediated ubiquitination that facilitate RNA processing. While all currently known cellular substrates of E1B55K/E4orf6 are degraded by the proteasome, we uncovered RNA-binding proteins (RBPs) as high-confidence substrates which are not decreased in overall abundance. We focused on two RBPs, RALY and hnRNP-C, which we confirm are ubiquitinated without degradation. Knockdown of RALY and hnRNP-C increased levels of viral RNA splicing, protein abundance, and progeny production during infection with E1B55K-deleted virus. Furthermore, infection with virus deleted for E1B55K resulted in increased interaction of hnRNP-C with viral RNA, and attenuation of viral RNA processing. These data suggest viral-mediated ubiquitination of RALY and hnRNP-C relieves a restriction on viral RNA processing, revealing an unexpected role for non-degradative ubiquitination in manipulation of cellular processes during virus infection.

## INTRODUCTION

Viruses have evolved mechanisms to alter cellular pathways to promote infection and inactivate host defenses. One way this can be achieved is through viral factors that redirect host post-translational protein modification such as ubiquitination, in order to regulate protein function and turnover. Viruses interface with the host ubiquitin system by encoding their own ubiquitin ligases, redirecting cellular ubiquitin ligases, or altering ubiquitin removal by deubiquitinating enzymes^1–3^. Ubiquitin can be employed as a signal for diverse outcomes, including proteasome-mediated degradation, protein localization, and regulating interactions with other proteins or nucleic acids^4–7^. This diversity of function makes hijacking the host ubiquitin machinery an attractive approach for viruses to manipulate multiple cellular pathways.

The nuclear-replicating Adenovirus (Ad) encodes two early proteins (E1B55K and E4orf6) which integrate into an existing host ubiquitin ligase complex containing Elongin B and C, Cullin5, and RBX1^8, 9^. The cellular ligase is recruited through E4orf6, and the E1B55K protein is involved in substrate recognition to redirect the ligase activity^9^. The importance of hijacking the host ubiquitin machinery for productive virus infection has been demonstrated using Ad deletion mutants or expression of dominant negative Cullin5, which all severely limit virus production^10–18^. Several cellular proteins have been identified as targets for proteasomal degradation mediated by the Ad serotype 5 (Ad5) E1B55K/E4orf6 complex, including MRE11, RAD50, NBS1, DNA Ligase IV, BLM, Integrin α3, and the tumor suppressor p53^8, 19–23^. Degradation of these proteins represses DNA damage signaling and apoptosis during infection^24–26^. However, the E1B55K/E4orf6 complex also stimulates export of viral late mRNAs and synthesis of viral late proteins^11–16^. Viral mutants defective for either E1B55K or E4orf6 exhibit reduced viral late RNA, late protein abundance, and progeny production but show little impact on early stages of virus infection^11–16^. The mutant virus phenotype was mapped to a nuclear step of viral late RNA processing. None of the known substrates fully explain these deficiencies, since mutant viruses still show lower late protein levels in cells deficient in p53 or lacking a functional DNA damage response^10–16^.

In this study, we used an unbiased global proteomics approach to identify new cellular substrates of ubiquitination mediated by the Ad5 E1B55K/E4orf6 complex. We used antibody-based di-glycine remnant enrichment combined with profiling by mass spectrometry (K-ε-GG)^27, 28^ to quantify changes in the cellular ubiquitinome induced upon expression of E1B55K and E4orf6. The K-ε-GG approach allows for direct identification of peptides modified as a result of E1B55K/E4orf6 expression. Furthermore, we examined the impact of ubiquitination on protein abundance by employing whole cell proteomics (WCP). This combined approach enabled us to identify many potential targets of the E1B55K/E4orf6 complex, and classify these proteins as predicted degraded or non-degraded substrates. Our analysis suggests that the E1B55K/E4orf6 complex can facilitate different types of ubiquitination, and reveals that the majority of cellular substrates are ubiquitinated without significant changes in their protein abundance. Among the cellular substrates predicted to be ubiquitinated without degradation, we found an enrichment for cellular RNA-binding proteins (RBPs). We further validated the importance of the highly ubiquitinated RBPs RALY and hnRNP-C as two host proteins modified by the virus to overcome restriction of viral late transcript production. We identify the first substrates that provide a mechanistic link between E1B55K/E4orf6-mediated ubiquitination and the known roles of the complex in Ad5 viral RNA processing. Furthermore, these studies highlight a viral approach to exploit ubiquitination without degradation as a strategy to manipulate host pathways.

## RESULTS

### Functional Ad E1B55K/E4orf6 complex is required for viral late RNA splicing

We hypothesized that ubiquitination mediated by the E1B55K/E4orf6 complex can either target cellular proteins for proteasomal degradation, as seen for all currently known substrates including MRE11, RAD50, and BLM^20, 23^, or could impact function without affecting protein abundance (**Fig. 1a**). We assessed the role of E1B55K/E4orf6-mediated ubiquitination on RNA processing and late protein accumulation by inactivating the complex through deletion of the E1B55K gene or chemical inhibition of Cullin5 ubiquitin ligase activity^29^. Infection with an E1B55K mutant virus (ΔE1B) resulted in decreased levels of viral late proteins (hexon, penton, fiber and protein VII) but had minimal impact on viral early protein production (DBP) when compared to wild-type (WT) Ad5 infection (**Fig. 1b; Supplementary Fig. 1a**). Cullin ubiquitin ligases require post-translational modification by the ubiquitin-like protein NEDD8 to form a functional ubiquitin ligase complex^30, 31^. We hypothesized that inhibition of the Cullin5 complex hijacked by E1B55K/E4orf6 would mimic ΔE1B virus infection. We used a small molecule inhibitor of the neddylation activating enzyme (NEDDi; MLN4924^29^) to block Cullin-mediated ubiquitination during infection. Inhibition of Cullin neddylation was confirmed by decreased abundance of the slower-migrating modified Cullin5 (**Fig. 1b**). Inhibition of the ubiquitin ligase activity of the viral E1B55K/E4orf6 complex was confirmed by blocking of MRE11 and BLM degradation. NEDDi treatment during WT Ad5 infection substantially decreased levels of viral late proteins (hexon, penton, fiber and protein VII) but only marginally decreased production of the viral early protein DBP (**Fig. 1b; Supplementary Fig. 1a**). Furthermore, NEDDi treatment did not further alter the late protein defect observed with E1B55K deletion (**Fig. 1b**). The increase of E1B55K levels upon NEDDi treatment is likely caused by inhibition of auto-ubiquitination, which is common among ubiquitin ligases^32^. We then assessed several steps of viral RNA processing during inhibition of ubiquitination or E1B55K deletion. We observed decreased accumulation of viral late mRNA for transcripts containing the major late promoter (MLP) and fiber gene during NEDDi treatment of WT Ad5 infection, similar to decreases detected with E1B55K deletion (**Fig. 1c**). These lower mRNA levels could be caused by defects in different steps of RNA processing: transcription, splicing, or decay. We assessed transcription and RNA turnover in WT and ΔE1B infection using 4sU-labeling of nascent RNA (**Supplementary Fig. 1b**). Our analysis revealed that deletion of E1B55K does not negatively impact transcription or RNA decay of viral early (E1A and E4) or viral late (MLP) RNA. Furthermore, we analyzed RNA decay by blocking transcription with Actinomycin D and measuring viral early (E1A) and late (MLP) RNA levels over a time course, comparing WT and ΔE1B infection (**Supplementary Fig. 1c**). This experiment confirmed that turnover of spliced viral RNA does not decrease in the absence of E1B55K. We used quantitative reverse transcription PCR (RT-qPCR) to determine the ratio of spliced:unspliced transcript as a measure for splicing efficiency (**Supplementary Fig. 1d**). This analysis revealed that both NEDDi treatment and E1B55K deletion decreased splicing efficiency of viral late transcripts (MLP and fiber), compared to untreated WT, without negatively impacting the early E1A transcript (**Fig. 1d**; **Supplementary Fig. 1e and 1f**). We also examined cytoplasmic RNA accumulation by fluorescence *in situ* hybridization (FISH) for fiber transcripts. This experiment demonstrated that less fiber RNA reaches the cytoplasm upon E1B55K deletion, which was recapitulated by NEDDi inhibition (**Fig. 1e**; **Supplementary Fig. 1g**). Failure to splice transcripts correctly causes retention in the nucleus and subsequent degradation of the unspliced RNA^33, 34^. Incorrect splicing could explain the observed RNA export defect and decrease in RNA levels observed for late viral transcripts. These data demonstrate that chemical inhibition of Cullin ligases recapitulates the effects of E1B55K deletion, highlighting that E1B55K/E4orf6-mediated ubiquitination of substrates is important for RNA splicing, RNA export, and protein production of viral late transcripts during Ad5 infection. None of the previously identified cellular substrates of E1B55K/E4orf6-mediated ubiquitination explain these phenotypes.

**Figure 1.**
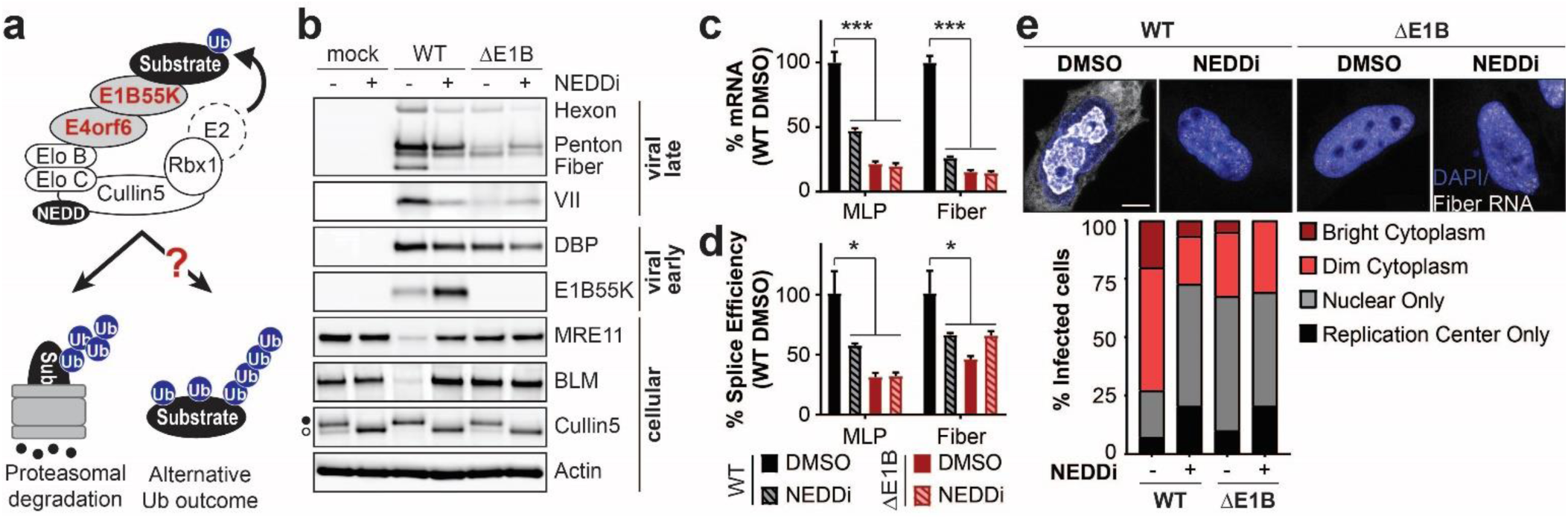
E1B55K deletion or inhibition of Cullin-mediated ubiquitination decreases adenovirus late RNA splicing and RNA processing overall. **a,** The E1B55K/E4orf6 complex redirects substrate recognition of the host Cullin5 ubiquitin ligase to target proteins for proteasomal degradation or induce alternative outcomes of ubiquitination. b-e, HeLa cells infected with wild-type (WT) or E1B55K-deleted (ΔE1B) Ad5 at multiplicity of infection (MOI) of 10. Cells were treated with either DMSO or NEDDi (neddylation inhibitor MLN4924) at 8 hours post-infection (hpi) and assayed at 24 hpi. b, Immunoblot analysis of viral and cellular protein abundance. The neddylated (●) and unmodified (○) forms of Cullin5 are indicated. Results are representative of three biological experiments. c, Bar graph representing spliced RNA levels of viral late transcripts for the major late promoter (MLP) and fiber by quantitative reverse transcription PCR (RT-qPCR). Shown is mean+s.d., n equals three biological experiments. d, Bar graph representing splicing efficiency as the ratio of spliced to unspliced transcripts of MLP and fiber relative to the WT DMSO control by RT-qPCR. Shown is mean+s.d., n equals three biological experiments. e, RNA FISH visualizing the localization of fiber transcripts (white) in relation to nuclear DNA stained with DAPI (blue) and quantification of observed pattern for > 50 HeLa cells. Scale bar 10 μm. Statistical significance was calculated using an unpaired, two-tailed Student’s t-test, * p < 0.05, *** p < 0.005.

### Proteomics reveals enrichment of RNA-binding proteins among cellular substrates of the E1B55K/E4orf6 complex

To identify cellular substrates of the Ad5 E1B55K/E4orf6 complex that could explain the RNA processing defect of the ΔE1B virus, we conducted global remnant profiling of the ubiquitinome (K-ε-GG) and associated whole cell proteome (WCP) over a time course of transduction of HeLa cells with viral vectors encoding Ad5 E1B55K and E4orf6^19, 35^ (**Fig. 2a**; **Supplementary Fig. 2**). Using non-replicating viral vectors allowed us to identify substrates specific to the activity of the viral E1B55K/E4orf6 complex outside the context of Ad5 infection. We assayed the degradation kinetics of known cellular substrates (BLM, MRE11, RAD50, and NBS1) by immunoblotting to determine when proteins were most likely to be modified but still detectable (**Supplementary Fig. 2a**). We subsequently performed K-ε-GG analysis for ubiquitin modification at 0, 6, 8, and 10 hours post-transduction (hpt), and WCP at 0 and 10 hpt for protein abundance^27, 28^ (**Fig. 2a**).

**Figure 2.**
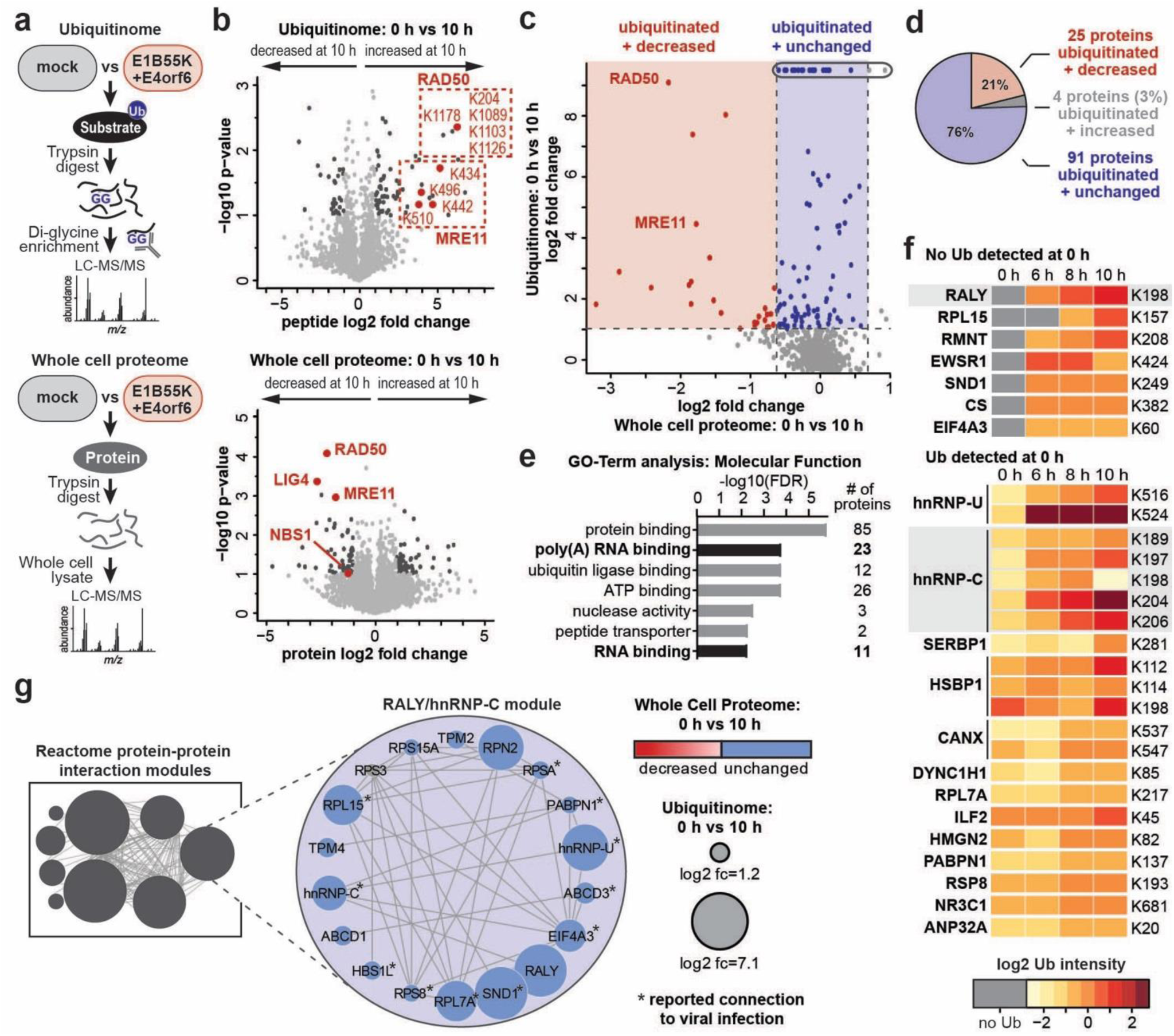
Unbiased proteomics reveals RNA-binding proteins among putative non-degraded substrates of the Ad E1B55K/E4orf6 complex. **a,** Proteomics workflow for identification of E1B55K/E4orf6 substrates. HeLa cells were transduced with recombinant Ad vectors expressing E1B55K and E4orf6 (MOI=10), and subjected to both di-glycine remnant profiling (K-ε-GG) to identify ubiquitinated lysine residues and whole cell proteomics to determine protein abundance. **b,** Volcano plots showing log2 fold-changes between 0 h and 10 h for ubiquitination (above) and protein abundance (below). For ubiquitination, individual peptides containing the modified lysine residues are normalized to protein abundance. Peptides and proteins with a fold change > ±s.e.m. and p-value < 0.05 are considered significantly changed and highlighted in dark grey. Ubiquitinated peptides and proteins corresponding to known E1B55K/E4orf6 substrates are highlighted in red. n equals three biological replicates. **c,** Scatter plot integrating changes in protein abundance (X-axis) and ubiquitination (Y-axis). Putative degraded substrates are shown in red (increased ubiquitination, decreased protein abundance), putative non-degraded substrates are shown in blue (increased ubiquitination, no significant change in protein abundance). Known degraded substrates MRE11 and RAD50 are indicated. Blue dots circled at the top indicate proteins that were only ubiquitinated upon expression of E1B55K/E4orf6 and were not detected as ubiquitinated in mock conditions. **d,** Bar graph representing gene ontology (GO) analysis of all predicted substrates by molecular functions. Categories containing RNA-binding proteins are highlighted. **e,** Predicted substrates that either decrease (red), increase (grey) or remain unchanged (blue) in their protein abundance during expression of E1B55K/E4orf6. **f,** Heat map of all ubiquitinated lysine residues within RNA-binding proteins with a normalized log2 abundance z-score > −0.5 and maximum log2 fold-change > 1 over the time course of E1B55K/E4orf6 transduction. The colors in the heat map correspond to the average z-score of the ubiquitination and are indicated in the accompanying scale. Highly ubiquitinated proteins RALY and hnRNP-C are highlighted. **g,** The Reactome-FI application in Cytoscape was utilized to generate a protein-protein interaction network in which nodes represent proteins and edges represent Reactome-based protein-protein interactions. Shown is the single module containing RALY and hnRNP-C. Node size corresponds to relative protein-based ubiquitination log2 fold change and node color corresponds to whole cell proteome log2 fold change following 10 h transduction of E1B55K/E4orf6. * denotes proteins that have a reported role during different viral infections.

Ubiquitin is covalently attached to its substrate and upon proteolytic cleavage with trypsin the C-terminal glycine residues of ubiquitin remain attached to the modified lysine residue (K-ε-GG). We enriched for peptides containing these di-glycine remnants using an antibody^27^ and identified modified peptides by mass spectrometry. We performed three replicates for each timepoint and identified a similar number of peptides in untransduced cells (2,050 peptides quantified in at least two replicates) and those transduced by E1B55K/E4orf6 at 6, 8, and 10 hours (2,132; 2,010; and 2,154 peptides respectively) (**Supplementary Fig. 2b; Supplementary Table 1**). The identified K-ε-GG peptides corresponded to 1,164 proteins overall. Changes in peptide modification were then normalized to changes in total protein abundance. Expression of E1B55K/E4orf6 induced a significant increase in ubiquitination (p < 0.05 and log2 fold-change > 1) for 39 peptides (**Fig. 2b**). Additionally, 51 peptides were ubiquitinated upon expression of E1B55K/E4orf6 but were not identified as ubiquitinated in untransduced cells, and therefore do not have a calculated fold-change or associated p-value. Peptides uniquely ubiquitinated during transduction are defined as those not quantified in any mock cell samples but found in 2-3 replicates from transduced cells. Since these unique peptides were not identified in mock conditions, they therefore do not have quantification values. The lack of quantification values precludes calculation of associated fold changes or p-values since both of these calculations require numerical values for both compared conditions. Therefore, in these cases, we used z-scores to assess abundance of ubiquitination during expression of E1B55K/E4orf6, and for downstream analysis to define the most highly ubiquitinated proteins. Peptides that exhibited increased or unique ubiquitination upon E1B55K/E4orf6 expression included known protein substrates MRE11 (4 peptides) and RAD50 (5 peptides).

A similar number of proteins were quantified in the WCP of untransduced cells (6,213 proteins identified in at least 2 replicates) and cells transduced by E1B55K/E4orf6 (6,241 proteins identified in at least 2 replicates in 10 hour timepoint) (**Supplementary Fig. 2c; Supplementary Table 1**). The WCP data show that E1B55K/E4orf6 expression induced significant changes in protein abundance, with 46 proteins significantly decreased at the 10 hour timepoint (log2 fold change <= −1 and p<0.05 or unique identification at 0 hour timepoint). Consistent with previous studies, we observed significant decreases for the known substrates MRE11, NBS1, RAD50, and LIG4 upon E1B55K/E4orf6 expression (**Fig. 2b**).

To compare K-ε-GG and WCP datasets, the peptide-level K-ε-GG data were transformed into protein-based K-ε-GG abundance changes by calculating the abundance-weighted average of the K-ε-GG peptide log2 fold-changes for all modified peptides detected for the respective protein. Resulting protein-based K-ε-GG log2 fold-changes were plotted against their associated WCP fold-changes (**Fig. 2c**). We implemented a threshold for protein-based K-ε-GG increase of > 2 fold and identified 120 host proteins as putative substrates of the Ad5 ubiquitin ligase. Proteins that were ubiquitinated and also decreased in abundance by more than 1 standard deviation (s.d.) from the mean proteome change were predicted to be degraded substrates of the E1B55K/E4orf6 complex (**Fig. 2c** and **2d**, red; **Supplementary Table 2**). The degraded substrates (25 proteins) include known targets MRE11 and RAD50. Conversely, proteins that were ubiquitinated and exhibited abundance changes within 1 s.d. of the mean WCP abundance change were predicted to be ubiquitinated as a result of the E1B55K/E4orf6 complex but not subsequently degraded (91 proteins) (**Fig. 2c** and **2d**, blue; **Supplementary Table 2**). These data provide the first evidence that the Ad5 E1B55K/E4orf6 complex facilitates non-degradative ubiquitination, and suggest that the majority of potential substrates of the viral complex fall into this category.

Gene ontology analysis of predicted substrates for the E1B55K/E4orf6 complex revealed significant enrichment of “poly(A) RNA binding” and “RNA-binding” GO annotations (**Fig. 2e; Supplementary Table 3**). Since E1B55K deletion has been shown to induce RNA processing defects, we focused on the 26 proteins included within the RNA-binding GO terms (**Supplementary Fig. 3a**). There were 7 RBPs predicted to be ubiquitinated only in the presence of E1B55K/E4orf6, of which RALY stands out as the RBP with the highest abundance of ubiquitination at 10 hpt. Additionally, hnRNP-C is an interaction partner of RALY which had the largest number of sites that increase in ubiquitination among RBPs (**Fig. 2f**; **Supplementary Fig. 3b**). We used the Reactome^36^ protein-protein interaction database to analyze interactions among all predicted substrates of the E1B55K/E4orf6 complex, and found RALY and hnRNP-C together in an interaction module with other RBPs (**Fig. 2g; Supplementary Fig. 4 and Table 3**). A literature search revealed that 11 of the 17 proteins in this module have reported association with viral infection (**Fig. 2g; Supplementary Table 4**). Both RALY and hnRNP-C are expressed at high levels in all tissues^37^ and are implicated in multiple steps of RNA processing, including RNA splicing and export^38–43^. Additionally, it has been reported that hnRNP-C binds to Ad transcripts encoding late proteins^44^. We therefore chose to further validate RALY and hnRNP-C as cellular substrates of the E1B55K/E4orf6 complex and to characterize their impact on Ad5 biology.

### RALY and hnRNP-C are ubiquitinated but not degraded upon E1B55K/E4orf6 expression

RALY and hnRNP-C are ∼43% homologous, with the highest homology (63%) in the coiled-coil (CC) domain, which contains all the lysine residues that show increased ubiquitination upon E1B55K/E4orf6 expression (**Fig. 3a**; **Supplementary Fig. 5**). The lysine residue of hnRNP-C that shows the highest increase in ubiquitination (K204) is analogous to the only detected ubiquitination site in RALY (K198). Since E1B55K is the substrate recognition component of the ligase assembled with the Ad complex, we examined interaction of E1B55K with the two host RBPs during Ad5 virus infection (**Fig. 3b**). We performed immunoprecipitation (IP) of E1B55K, RALY and hnRNP-C for mock, Ad5 WT, and ΔE1B infection conditions followed by immunoblotting for viral and host proteins. IP of E1B55K isolated RALY and hnRNP-C from cells infected with WT virus but not the ΔE1B mutant (**Fig. 3b**). In the reciprocal experiment, E1B55K was detected upon IP of either RALY or hnRNP-C during WT virus infection, confirming interaction between the E1B55K/E4orf6 complex and the two host RBPs (**Fig. 3b**). The cellular hnRNP-C and RALY proteins interact in reciprocal IPs, as reported previously^45^, and this association was not impacted by virus infection. To confirm ubiquitination of RALY and hnRNP-C induced by viral proteins, we expressed Flag-tagged RALY or hnRNP-C, HA-tagged ubiquitin, and E4orf6 by transfection of HEK293 cells (this cell line contains a genomic integration of Ad5 E1B55K^46^). IP for the HA epitope and immunoblotting for Flag revealed an increase in high molecular weight ubiquitin-complexes of RALY and hnRNP-C in the presence of E4orf6 (**Fig. 3c; Supplementary Fig. 6a**). hnRNP-C2, an alternative isoform of hnRNP-C, is also ubiquitinated in the presence of the E1B55K/E4orf6 complex (**Supplementary Fig. 6b**). The overall stoichiometry for ubiquitination is low relative to the total protein abundance of RALY and hnRNP-C, as is the case with many post-translational protein modifications. To demonstrate that activated Cullin complexes are involved in ubiquitination of RALY and hnRNP-C by the Ad5 complex, we performed experiments with NEDDi treatment. The elevated ubiquitination of RALY and hnRNP-C detected by expression of E4orf6 and HA-ubiquitin was decreased upon NEDDi treatment (**Fig. 3d; Supplementary Fig. 6c**). Inhibition of NEDDylation appeared to reduce endogenous ubiquitination of hnRNP-C but not RALY. These data suggest that a Cullin ubiquitin ligase may ubiquitinate hnRNP-C but not RALY in the absence of the viral proteins, consistent with the K-ε-GG data where hnRNP-C modification was increased from the mock condition but RALY was uniquely modified during infection. We also verified that hnRNP-C was ubiquitinated during infection with Ad5 WT but not with the ΔE1B mutant (**Fig. 3e**). The RALY antibody quality precluded our ability to detect endogenous protein in this assay. Our WCP analysis showed that RALY and hnRNP-C are not decreased in abundance during infection (**Supplementary Fig. 3a**). Lack of degradation was confirmed by immunoblot analysis which showed stable abundance of RALY and hnRNP-C protein levels over a time course of Ad5 WT infection or transduction with E1B55K/E4orf6 vectors (**Fig. 3f**; **Supplementary Fig. 6d**). RALY and hnRNP-C levels were also stable during transduction of A549 and U2OS cells with E1B55K/E4orf6 vectors, as well as transfection of HEK293 cells with an E4orf6 expression vector (**Supplementary Fig. 6e**). RALY and hnRNP-C protein levels remained relatively stable during infection in the presence of cycloheximide, further supporting that turnover is not increased by infection (**Fig. 3g**). Finally, mRNA levels for RALY and hnRNP-C as measured by RT-qPCR, remain stable during a time course of Ad WT infection (**Supplementary Fig. 6f**). We hypothesize that the E1B55K/E4orf6 complex facilitates ubiquitination that induces degradative and non-degradative outcomes, depending on the substrate. To test this hypothesis, we investigated differences in ubiquitination of MRE11, RAD50, RALY, and hnRNP-C mediated by the E1B55K/E4orf6 complex. Proteasome inhibition by drugs such as MG132 leads to accumulation of ubiquitinated proteins that would otherwise be degraded. Ubiquitination assays were performed by transfection of HEK293 cells with and without MG132-mediated proteasome inhibition (**Fig. 3h**). Expression of E4orf6 increased ubiquitination of MRE11 which was further increased by proteasome inhibition, consistent with MRE11 being a known degraded substrate of the viral complex. In contrast, expression of E4orf6 increased ubiquitination of RALY and hnRNP-C but there was no further increase upon treatment with MG132. The fact that MG132 treatment did not alter ubiquitination of RALY and hnRNP-C suggests that ubiquitination of these substrates does not result in degradation by the proteasome. Since the effect of proteasomal inhibition varies between substrates of the E1B55K/E4orf6 complex, we examined the ubiquitin chains attached to RALY and hnRNP-C as compared to MRE11 and RAD50. The ubiquitin linkage most commonly associated with proteasomal degradation is K48. To determine whether K48-linked ubiquitin is attached to MRE11, RAD50, RALY, or hnRNP-C we performed native IPs of HA-ubiquitin, expressed in HEK293 cells together with E4orf6, and then compared the degree of ubiquitination after treatment with deubiquitinating enzymes (DUBs) that cleave either all ubiquitin linkages (DUB^Pan^) or only K48-linked ubiquitin chains (DUB^K48^)^47^ (**Fig. 3i; Supplementary Fig. 6g**). MRE11 and RAD50 showed a clear decrease of high molecular weight ubiquitin chains upon treatment with both DUBs, indicating that K48-linked ubiquitin is attached to these substrates to induce proteasomal degradation. In contrast, ubiquitination of RALY and hnRNP-C decreased with the DUB^Pan^ but not the more specific DUB^K48^. This suggests that RALY and hnRNP-C are substrates for non-K48 linked ubiquitination, distinct from the K48-linked ubiquitin chains on degraded substrates MRE11 and RAD50. Our data support a non-degradative role for ubiquitination of hnRNP-C and RALY, although it is possible that degradation occurs within a sub-population too small to distinguish by this global analysis. Together, these data validate RALY and hnRNP-C as the first non-degraded cellular substrates identified for the E1B55K/E4orf6 Ad5 ligase complex.

**Figure 3.**
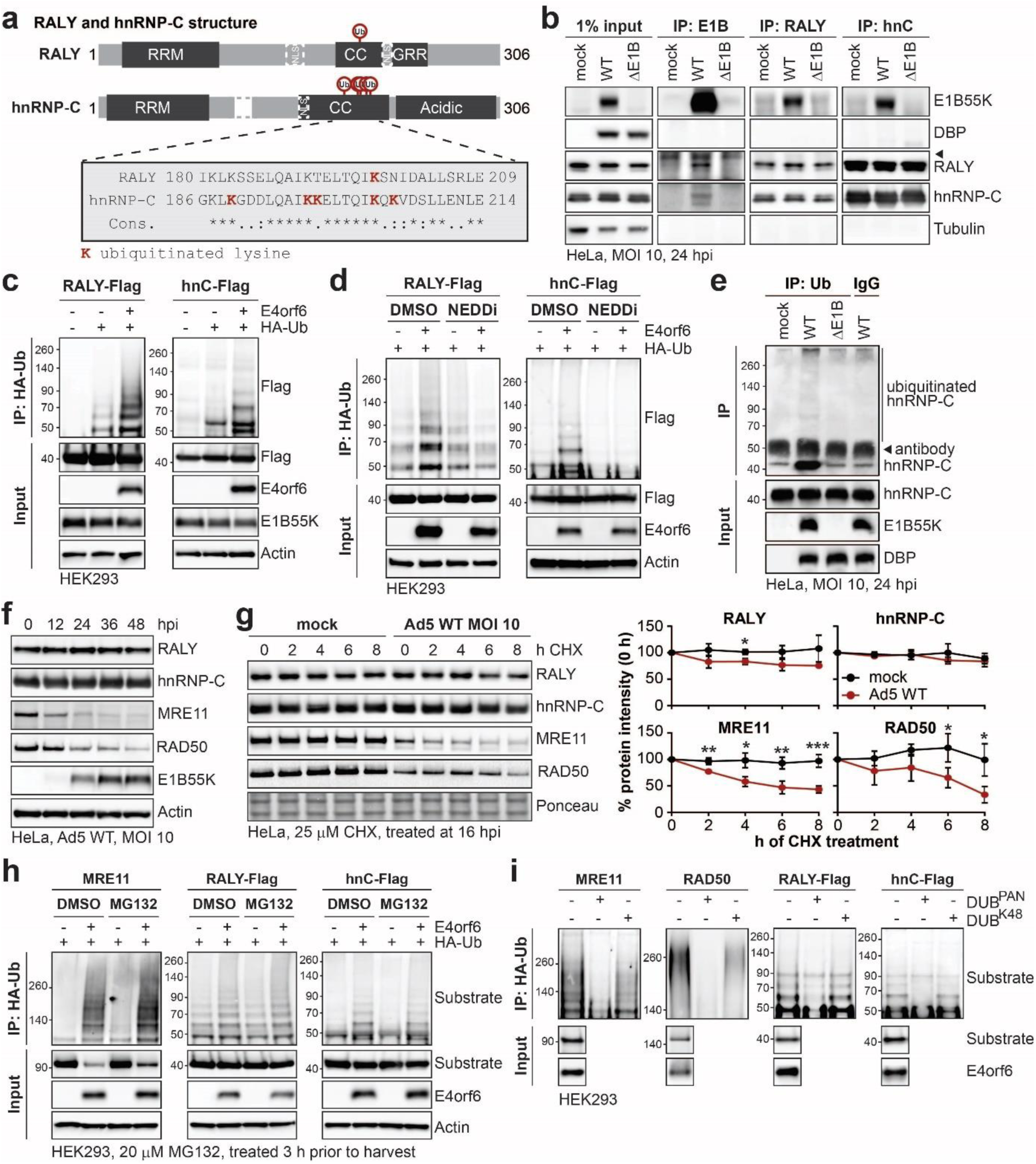
RALY and hnRNP-C are non-degraded substrates of the Ad E1B55K/E4orf6 complex ubiquitin ligase activity. **a,** Domain structure of RALY and hnRNP-C. RRM = RNA recognition motif, NLS = nuclear localization sequence, CC = coiled-coil region, GRR = glycine rich region. CC region shown below contains all the lysine residues with increased ubiquitination upon E1B55K/E4orf6 expression highlighted in red. **b,** Immunoblot analysis of E1B55K, RALY, and hnRNP-C immunoprecipitations (IP) probing for pull-down of viral and cellular proteins during mock, WT and ΔE1B infection of HeLa cells at MOI of 10 for 24 hpi. The viral protein DBP and cellular protein Tubulin served as negative controls and were not isolated with any condition. denotes the signal of the antibody heavy chain. **c,** HEK293 cells transfected with the indicated constructs for 24 h followed by denaturing IP with HA antibody and immunoblot analysis of RALY-Flag or hnRNP-C-Flag. **d,** HEK293 cells transfected with the indicated constructs for 24 h and treated with DMSO or NEDDi 6 h prior to harvest followed by denaturing IP with HA antibody and immunoblot analysis of RALY-Flag or hnRNP-C-Flag. **e,** Immunoblot of denaturing hnRNP-C IP probing for ubiquitin during mock, WT and ΔE1B infections at MOI of 10 for 24 h. indicates non-specific signal of the antibody heavy chain. **f,** Immunoblot analysis of protein levels over a time course of Ad5 WT infection (MOI=10) of HeLa cells. **g,** Immunoblot analysis and quantification of RALY, hnRNP-C, MRE11, and RAD50 over a time course of cycloheximide (CHX) treatment of mock or Ad5 WT infected HeLa cells. Quantification showing mean+s.d. of three biological replicates. Statistical significance was calculated using an unpaired, two-tailed Student’s t-test, * p < 0.05, ** p < 0.01, *** p < 0.005. **h,** HEK293 cells transfected with the indicated constructs for 24 h and treated with DMSO or proteasome inhibitor MG132 3 h prior to harvest followed by denaturing IP with HA antibody and immunoblot analysis of MRE11, RALY-Flag, and hnRNP-C-Flag. **i,** HEK293 cells transfected with the indicated constructs for 24 h followed by denaturing IP with HA antibody, treatment with the indicated deubiquitinating enzymes (DUBs) and immunoblot analysis of MRE11, RAD50, RALY-Flag, and hnRNP-C-Flag. All immunoblots are representative of at least three biological replicates. Size markers in kDa are shown for ubiquitination immunoblots.

### RALY and hnRNP-C are detrimental for viral late RNA processing

To determine whether RALY and hnRNP-C impact Ad infection, we used siRNA to knockdown these host proteins in HeLa and primary-like HBEC3-KT cells, and then infected with WT Ad5 and ΔE1B viruses (**Fig. 4**). Although RALY and hnRNP-C are not degraded during infection, this approach allowed us to determine whether these RBPs are beneficial or detrimental to virus infection. Knockdown of RALY and hnRNP-C did not affect viral protein levels during WT Ad5 infection, suggesting that in the context of infection with a fully competent virus their presence does not have a significant impact. Infection with ΔE1B virus generated reduced viral late protein levels as compared to WT Ad5 (**Fig. 4a**). Depletion of RALY and hnRNP-C rescued this viral late protein defect almost to the level observed in WT Ad5 (**Fig. 4a**; **Supplementary Fig. 7a and 7b**). We examined whether knockdown of RALY and hnRNP-C also affects progeny production of the ΔE1B virus (**Fig. 4b**). There was no difference between WT Ad5 and ΔE1B at 8 hours post-infection (hpi), before production of new infectious virions, confirming comparable virus input and entry. By 24 hpi the ΔE1B virus produced > 100-fold fewer viral particles than WT Ad5. Knockdown of RALY and hnRNP-C had no effect on WT Ad5, but significantly increased progeny production for the mutant virus (**Fig. 4b**). Similar rescue of the ΔE1B virus was observed with RALY and hnRNP-C knockdown prior to infection in HBEC3-KT cells (**Fig. 4b**). These data suggest that RALY and hnRNP-C are detrimental to Ad infection and that E1B55K/E4orf6-mediated ubiquitination relieves their restriction on virus production. Since RALY and hnRNP-C are involved in RNA splicing and export, we hypothesized that their depletion selectively increases late RNA processing without affecting DNA replication and viral early RNAs. We therefore examined viral DNA replication by quantifying genome accumulation using qPCR (**Fig. 4c**). There was a modest decrease (2-fold) in DNA replication for the ΔE1B virus as compared to WT Ad5, in agreement with prior reports^48^. Viral DNA accumulation for both WT Ad5 and ΔE1B was not significantly affected by depletion of RALY and hnRNP-C (**Fig. 4c**), confirming that their effects are mediated at a step after viral genome replication. We then quantified RNA levels of both viral early (E1A) and late (MLP and fiber) transcripts (**Fig. 4d**). Levels of late but not early transcripts decreased upon infection with ΔE1B virus, which shows qualitative correlation with the decrease in late proteins shown in in **Fig. 1b**. Depletion of RALY and hnRNP-C rescued mRNA levels for MLP and fiber at both 18 hpi and 24 hpi during infection with ΔE1B virus, to levels observed in WT Ad5 (**Fig. 4d**) without impacting the E1A transcript (**Supplementary Fig. 7c**). Splicing efficiency of MLP and fiber was reduced in the ΔE1B virus and was rescued to WT Ad5 levels upon knockdown of RALY and hnRNP-C (**Fig. 4e**; **Supplementary Fig. 7c**). We also used FISH to examine the effect of RALY and hnRNP-C depletion on export of fiber mRNA into the cytoplasm. siRNA treatment increased the amount of cytoplasmic fiber RNA visible in ΔE1B infection, while not impacting WT Ad5 (**Fig. 4f**). Depletion of either RALY or hnRNP-C by itself increased viral late protein, RNA levels, and splicing efficiency of the mutant virus, with hnRNP-C knockdown having a more dramatic effect than RALY knockdown (**Supplementary Fig. 7e-g**). To connect the impact of RALY and hnRNP-C depletion on late stages of Ad infection with Cullin-dependent ubiquitination by the E1B55K/E4orf6 complex, we combined siRNA-mediated knockdown with NEDDi treatment during WT Ad5 infection. The NEDDi treatment decreased viral late RNA levels, splicing efficiency, and protein production (**Fig. 4g-i**). Knockdown of RALY and hnRNP-C completely rescued the defect caused by inhibition of Cullin function without impacting viral early proteins or RNA (**Fig. 4g-i**; **Supplementary Fig. 7h-j**). These data suggest that RALY and hnRNP-C are detrimental to the late stages of Ad5 infection and that ubiquitination or depletion can overcome this defect.

**Figure 4.**
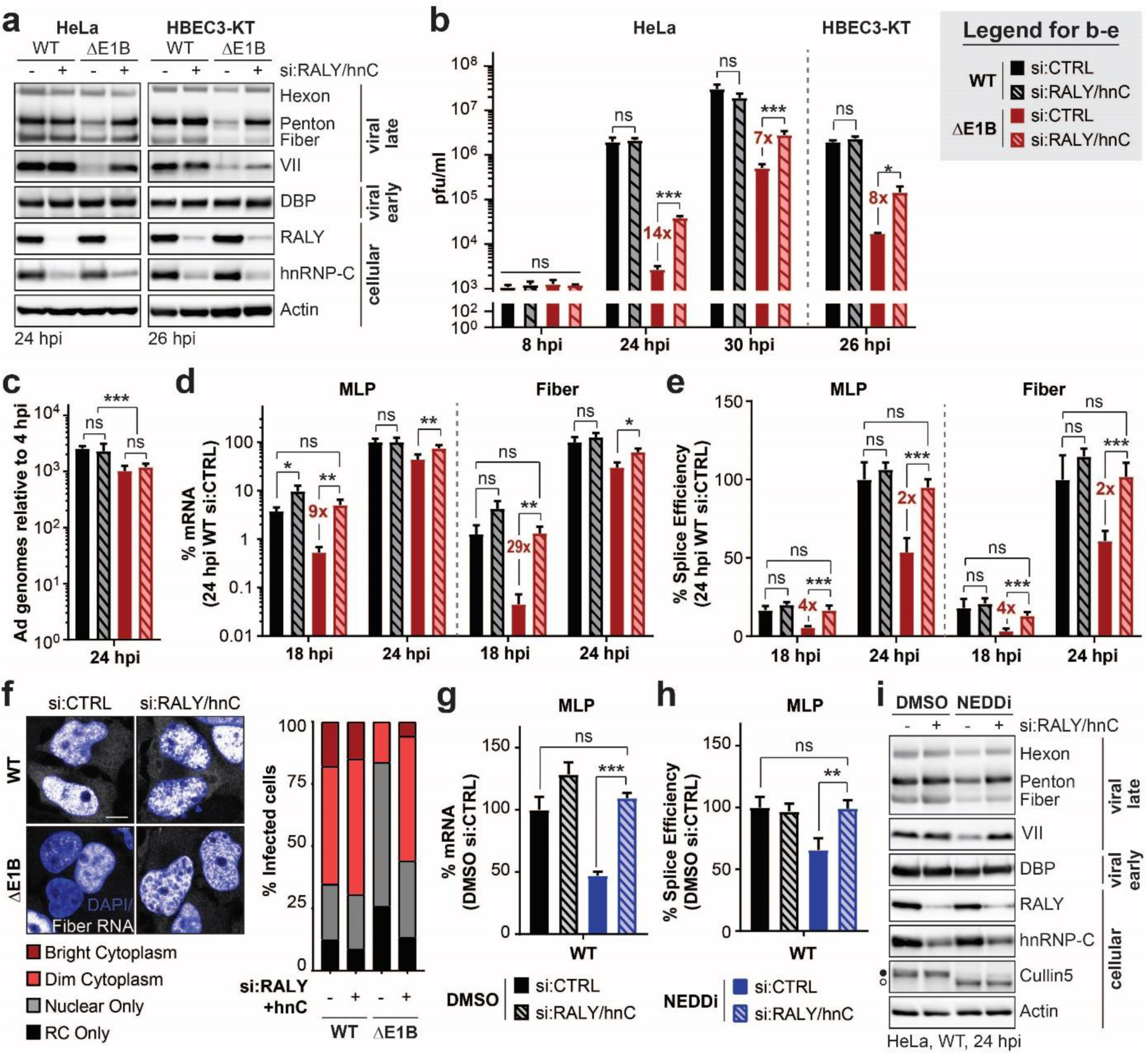
Knockdown of RALY and hnRNP-C rescues the RNA processing defect caused by the absence of a functional E1B55K/E4orf6 complex. **a-f,** HeLa cells or HBEC3-KT (only a,b) transfected with control (si:CTRL) or RALY and hnRNP-C (si:RALY/hnC) siRNA 24 h prior to infection with Ad5 WT or ΔE1B (MOI 10), harvested at respective time points. **a,** Immunoblot analysis of viral and cellular protein levels. **b,** Bar graph representing plaque assays for viral progeny. pfu = plaque forming units. **c,** Bar graph representing qPCR of viral genomes normalized to input. **d,** Bar graph representing spliced RNA levels of viral late transcripts MLP and fiber measured by RT-qPCR. **e,** Bar graph representing splicing efficiency as defined as the ratio of spliced to unspliced transcripts of MLP and fiber measured by RT-qPCR. **f,** RNA FISH visualizing the localization of fiber transcripts (white) in relation to nuclear DNA stained with DAPI (blue) and quantification of observed pattern for > 100 HeLa cells. RC – replication center. Scale bar 10 μm. **g-i.** HeLa cells transfected with control (si:CTRL) or RALY and hnRNP-C (si:RALY/hnC) siRNA 24 h prior to infection with Ad5 WT (MOI=10), treated with either DMSO or NEDDi at 8 hpi and processed at 24 hpi. **g,** Bar graph representing spliced RNA levels of MLP measured by RT-qPCR. **h,** Bar graph representing splicing efficiency as defined as the ratio of spliced to unspliced transcripts of MLP measured by RT-qPCR. **i** Immunoblot analysis of viral and cellular protein levels, with neddylated (●) and unmodified (○) forms of Cullin5 indicated. All immunoblots are representative of at least three biological experiments. All graphs show the mean+s.d. with n equals three biological replicates. Statistical significance was calculated using an unpaired, two-tailed Student’s t-test, * p < 0.05, ** p < 0.01, *** p < 0.005.

### Infection causes global changes to hnRNP-C RNA binding

Our data suggest that viral-mediated ubiquitination of RALY and hnRNP-C relieves a restriction on viral late RNA processing without the need for proteasomal degradation. Non-degradative ubiquitination has been reported to alter protein localization, for example by obscuring nuclear localization sequences and preventing nuclear import^49^. We examined localization of RALY and hnRNP-C by immunofluorescence (IF) in untreated HeLa cells and during infection with either WT Ad5 or ΔE1B virus (**Supplementary Fig. 8a**). Both RALY and hnRNP-C showed a diffuse nuclear pattern in uninfected HeLa cells, in accordance with the reported localization of both proteins^37^. Upon infection, both proteins were excluded from viral replication centers marked by DBP or USP7 in a pattern that matches viral RNA and other RBPs^50, 51^. However, there was no obvious difference in localization between WT Ad5 and ΔE1B infection, suggesting that viral-induced ubiquitination does not specifically change their cellular localization. Since both RALY and hnRNP-C are ubiquitinated within the coiled-coil domain that is involved in multimerization and protein-RNA interaction (**Fig. 3a**), we examined whether overall protein complex formation is affected by treating HeLa cells with disuccinimidyl suberate (DSS) at various concentrations during mock, WT Ad5 or ΔE1B infection (**Supplementary Fig. 8b**). DSS is a cell-permeable crosslinker that forms stable amide bonds between lysine residues in close proximity (less than 11.4 Å), crosslinking protein complexes. DSS treatment caused a mobility shift of hnRNP-C and RALY, consistent with previous reports of multimerization^52^. During WT Ad5 or ΔE1B infections these patterns did not change, suggesting that viral-induced ubiquitination does not significantly affect overall protein complex formation of hnRNP-C or RALY. Next, we employed targeted proteomic identification of RNA-binding regions (RBR-ID)^53, 54^ for hnRNP-C (**Supplementary Fig. 8c; Supplementary Table 5**). We compared in triplicate SILAC-labeled^55^ HeLa cells that were uninfected or infected with Ad5 WT or ΔE1B. At 24 hpi we performed 4sU-mediated protein-RNA photo-crosslinking of heavy-labeled cells, followed by hnRNP-C IP, nuclease treatment and proteolytic cleavage. Peptides in the crosslinked condition that bound RNA will retain RNA adducts, which causes a mass shift and loss of signal when compared to peptides from non-crosslinked conditions. Signal loss thus identifies which regions of a protein had direct contact with RNA *in vivo*. In addition, the non-crosslinked data give insight into the hnRNP-C interactome and potential changes upon Ad infection. In mock conditions, the most dramatic loss of signal was detected at the RNA-recognition motif (RRM) within hnRNP-C (**Fig. 5a**). The RRM is the most well-characterized RNA-binding domain in hnRNP-C and provides specificity for the poly-U motif identified as the preferred binding site^56^. Surprisingly, upon both WT and ΔE1B infection the RRM interaction with RNA was dramatically decreased, while RNA interactions within the coiled-coil domain increased (**Fig. 5a**). Approximately ∼20 amino acids downstream of the ubiquitination sites, we detected an RNA binding peak in mock samples that was decreased in WT Ad5 but increased in the mutant virus (**Fig. 5a**). This observation highlights a region of hnRNP-C potentially impacted by Ad-mediated ubiquitination. Since these two RBPs interact strongly, we were also able to analyze RNA binding for RALY from the hnRNP-C IP. Similar to hnRNP-C, we saw infection-mediated changes in the interaction of the RRM with RNA, and potential ubiquitin-mediated differences between WT and ΔE1B infection close to the ubiquitination site (**Supplementary Fig. 8d**). In contrast to the large differences observed for the RNA-binding analysis, we only observed minimal differences in the hnRNP-C interactome when comparing mock, WT, and ΔE1B infection (**Supplementary Fig. 8e; Supplementary Tables 6**). A global comparison of the interacting protein abundances across conditions revealed a Pearson correlation coefficient of >0.8. In addition, the 20 most abundant hnRNP-C interactors did not show marked differences in interaction abundance between mock, WT, and ΔE1B infection, with the only exception being viral proteins absent in mock (**Supplementary Fig. 8f**). In summary, RBR-ID revealed major changes to RNA binding of hnRNP-C during infection and a potential ubiquitin-mediated difference between Ad5 WT and ΔE1B infections.

**Figure 5.**
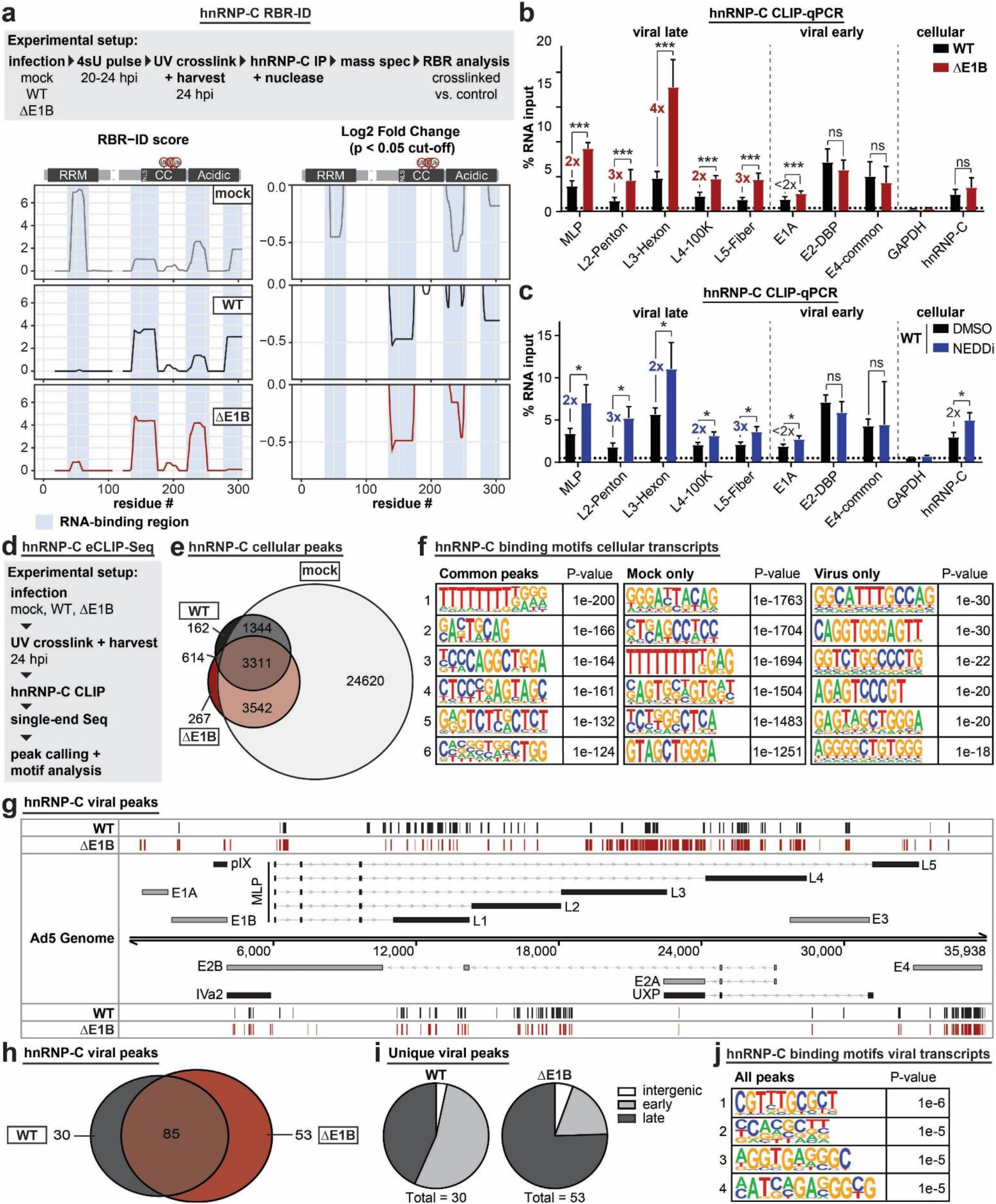
The interaction of hnRNP-C with viral late RNA increases in the absence of a functional E1B55K/E4orf6 complex. **a,** RBR-ID (RNA-binding region identification) for hnRNP-C comparing mock (grey), Ad5 WT (black), and ΔE1B (red) at 24 hpi with MOI 10. Shown are the experimental setup (**above**), smoothed residue-level RBR-ID score plotted along the primary sequence (**left**), and smoothed residue-level fold-change between crosslinked and control conditions with a significance threshold of p < 0.05 (**right**). hnRNP-C domain structure with ubiquitination sites is shown above graphs. RNA-binding regions are highlighted in blue shaded area. **b,** HeLa cells infected with either WT Ad5 or ΔE1B (MOI=10), UV-crosslinked and harvested at 24 hpi, subjected to hnRNP-C CLIP and RT-qPCR for viral early and late transcripts. GAPDH is a cellular negative control. hnRNP-C is a cellular positive control. **c,** HeLa cells infected with WT Ad5 (MOI=10), treated with either DMSO or NEDDi at 8 hpi, UV-crosslinked and harvested at 24 hpi, subjected to hnRNP-C CLIP and RT-qPCR for viral early and late transcripts. GAPDH is a cellular negative control. hnRNP-C is a cellular positive control. Graphs show mean+s.d, n equals six (b) or three (c) biological replicates. Statistical significance was calculated using an unpaired, two-tailed Student’s t-test, * p < 0.05, ** p < 0.01, *** p < 0.5. **d,** Experimental setup for hnRNP-C eCLIP-Seq. **e,** Venn diagram showing the overlap of hnRNP-C peaks called in host transcripts for mock (grey), Ad5 WT (black), and ΔE1B (red). **f,** Top six motifs identified for hnRNP-C binding sites in host transcripts present in all 3 conditions (left), mock only (middle), and virus only (right, WT only + ΔE1B only + WT and ΔE1B). **g,** hnRNP-C peaks called for Ad transcripts in Ad5 WT (black) and ΔE1B (red) infection on the forward strand (above) and reverse strand (below). The simplified schematic of the viral transcriptome shows forward facing transcription units above the genome and reverse facing transcription units below. Viral genes are color-coded to denote early (grey) and late (black) transcription units. Lines with arrows denote introns and bars are exonic regions. **h,** Venn diagram showing the overlap of hnRNP-C peaks called in viral transcripts for Ad5 WT (black) and ΔE1B (red). **i.** Pie charts of unique peaks for WT and ΔE1B showing the location in intergenic, early, or late transcription units. **j,** Top four motifs identified for hnRNP-C binding sites in viral transcripts present in any of the conditions.

### Interaction of hnRNP-C and RALY with viral late RNA is increased when Ad-mediated ubiquitination is disrupted

RBR-ID identifies sites of RNA binding within a protein sequence but does not identify the RNA sequence that is bound. To determine the impact of Ad-mediated ubiquitination on interaction of hnRNP-C with viral RNA we performed crosslinking-immunoprecipitation (CLIP) followed by RT-qPCR for viral and cellular transcripts (**Supplementary Fig. 9a and 9b**). The hnRNP-C transcript itself served as a positive control for immunoprecipitation, while GAPDH RNA was a negative control^57^. All viral late transcripts were detected above background under WT Ad5 conditions, however, there was a 2 to 4-fold increase in the amount of late RNA detected during ΔE1B infection. There was however no dramatic difference in the level of early RNAs detected between WT Ad5 and mutant virus. This indicates that viral-induced ubiquitination of hnRNP-C specifically decreases the interaction with viral late transcripts. Since the overall stoichiometry for ubiquitination is low relative to the total protein abundance, this could indicate that ubiquitination either has a dominant negative impact on the overall protein pool or that the effect is localized. This approach showed linearity over a ten-fold dilution of input material, and displayed the same trend of increased binding to viral late RNA upon ΔE1B infection (**Supplementary Fig. 9c**). In contrast, hnRNP-C CLIP-qPCR without UV-crosslinking or a CLIP with an IgG control precipitated minimal RNA (**Supplementary Fig. 9d and 9e**). Commercially available antibodies for RALY were not suitable for this technique. Therefore, we created an inducible RALY-Flag cell line and performed CLIP-qPCR by Flag immunoprecipitation. This demonstrated that similarly to hnRNP-C, RALY interacts more with viral late RNA in ΔE1B infections, while binding to early RNA is unchanged or even decreased (**Supplementary Fig. 9f**). To support the idea that differences in hnRNP-C interaction with viral RNA are caused by ubiquitination, we repeated the hnRNP-C CLIP-qPCR with inhibition of Cullin-dependent ubiquitination during WT Ad5 infection (**Fig. 5c, Supplementary Fig. 9g**). Following the trend with ΔE1B infection, the interaction of hnRNP-C with viral late transcripts increased at least 2-fold upon treatment with NEDDi, while there were only minor differences for viral early and cellular transcripts. This experiment reinforces that hnRNP-C interaction with viral late RNAs increases in the absence of the functional viral ubiquitin ligase complex.

To determine whether hnRNP-C binding changes in an ubiquitin-mediated manner during infection, we performed a global analysis of hnRNP-C binding sites comparing mock, Ad5 WT, and ΔE1B infection using enhanced CLIP followed by sequencing (eCLIP-Seq^58^) (**Fig. 5d**; **Supplementary Fig. 9h and 9i**). We observed a dramatic reduction of hnRNP-C binding to host RNAs upon infection (**Fig. 5e; Supplementary Fig. 9j**). More than 24,000 peaks were unique to mock condition, with only ∼3,000 identified in all 3 conditions. Analysis of binding motifs revealed the known poly-U/poly-T binding motif of hnRNP-C among common and mock specific peaks (**Fig. 5f**). We also detected hnRNP-C peaks in host RNA that were only identified during WT and ΔE1B infection, suggesting a potential role for ubiquitination in manipulating binding of hnRNP-C to cellular transcripts (**Supplementary Fig. 9k**). The hnRNP-C poly-U motif was lacking in these virus-specific peaks, supporting the RBR-ID data which show decrease of RNA binding for the hnRNP-C RRM upon infection (**Fig. 5a**). We also analyzed hnRNP-C binding sites on viral transcripts (**Fig. 5g** and **5h**). The number and location of hnRNP-C peaks were different between WT and ΔE1B infection. Analyzing peaks unique to WT or mutant virus revealed that deletion of E1B55K increased hnRNP-C binding mainly in viral late transcripts (**Fig. 5i**). Differences were especially pronounced in the L3-L5 region of the major late transcription unit (**Fig. 5g**), which encodes viral hexon protein. These results were consistent with our CLIP-qPCR data (**Fig. 5b**). In addition, there are several hnRNP-C binding sites in viral late RNA regions such as MLP and fiber that are unique to ΔE1B infection (**Fig. 5g**). Finally, we analyzed motifs present at hnRNP-C binding sites on viral transcripts. We saw no evidence of the canonical hnRNP-C poly-U motif observed on host transcripts (**Fig. 5j**). The most prominent motif for infection-specific hnRNP-C binding sites within both host and viral transcripts is very similar, suggesting a potential shift to a new hnRNP-C recognition motif caused by Ad5 infection. In summary, the RBR-ID and eCLIP-Seq data highlight major changes in hnRNP-C interaction with RNA caused by infection. Together, these results support a mode in which ubiquitination of hnRNP-C and RALY induced by the E1B55K/E4orf6 complex leads to reduced interaction of these host RBPs with viral late RNA, thereby overcoming a detrimental effect on viral RNA processing (**Fig. 6**).

**Figure 6.**
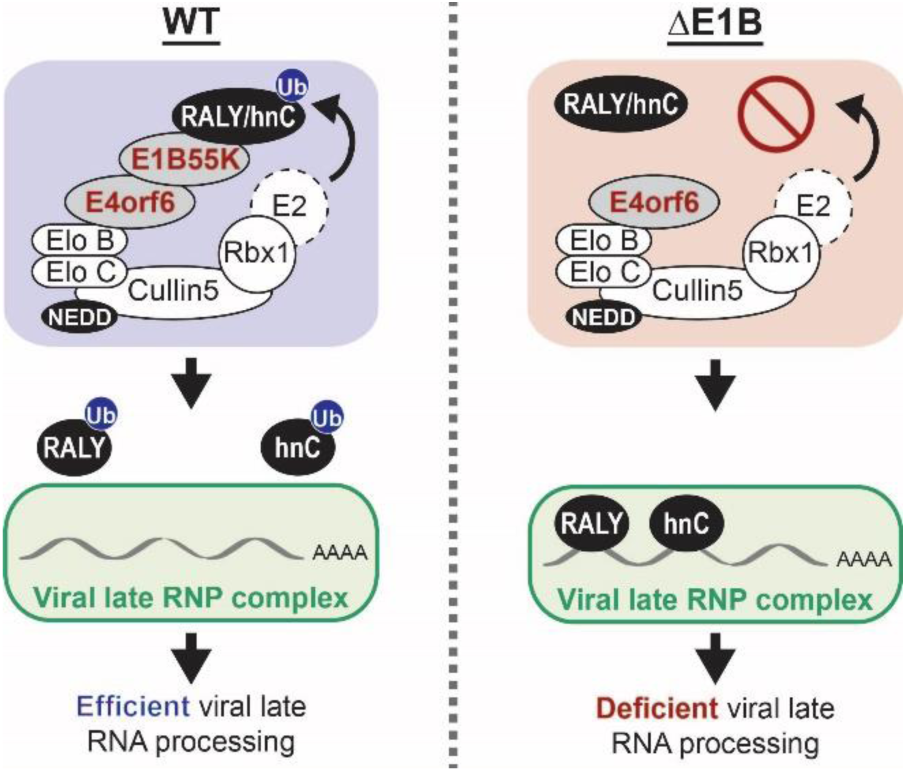
Non-degradative ubiquitination of RNA-binding proteins promotes efficient adenoviral RNA processing. During wild-type (WT) Ad5 infection the E1B55K/E4orf6 complex induces ubiquitination of RNA-binding proteins RALY and hnRNP-C to facilitate efficient viral late RNA processing. Ubiquitination regulates interaction of these host proteins with viral RNA to facilitate viral infection. In the absence of the E1B55K/E4orf6 complex ubiquitin ligase activity, the RBPs bind more to viral late mRNAs and limit RNA processing and protein production. RNP – ribonucleoprotein.

## DISCUSSION

Viruses commonly adapt cellular regulatory mechanisms towards efficient viral production. The E1B55K/E4orf6 complex is known to interact with the cellular Cullin5 ubiquitin ligase to redirect ubiquitination and to stimulate viral late mRNA nuclear export and late protein synthesis. Prior studies identified binding partners of the complex and a limited number of substrates^8, 9, 19–23, 59–62^, however, these studies did not enrich for proteins specifically ubiquitinated by the E1B55K/E4orf6 complex or explicitly link potential cellular substrates to effects on viral RNA processing. Here we employed a systematic proteomics approach to identify cellular ubiquitination substrates of the viral E1B55K/E4orf6 complex by combining quantification and analysis of the ubiquitinome and the associated whole cell proteome upon expression of E1B55K/E4orf6. We identified 119 potential substrates, with specific enrichment of RBPs that may be involved in viral RNA processing. In addition to RNA processing, functional analysis of the predicted substrates highlighted other host pathways that may be manipulated by Ad5-mediated ubiquitination: ubiquitin machinery and de-ubiquitinating enzymes, antigen presentation, protein folding, cellular transport, and cell signaling (**Supplementary Fig. 4a**). We focused on two of the most highly ubiquitinated RBPs, RALY and hnRNP-C, which we demonstrated to be the first non-degraded ubiquitination substrates of the Ad5 complex. We demonstrated differential interaction of hnRNP-C and RALY with viral late transcripts in the presence of an active E1B55K/E4orf6 complex, supporting a model in which RALY and hnRNP-C ubiquitination results in altered binding to viral late ribonucleoprotein (RNP) complexes, to promote efficient processing of late RNA (**Fig. 6**). Since hnRNP-C has reported roles in alternative splicing^38, 40, 41^, we propose that ubiquitination by the Ad5-induced complex results in exclusion from viral RNP complexes to promote splicing of late viral RNAs. Substrates of the E1B55K/E4orf6 complex can vary across human Ad serotypes, although some target proteins fall within the same cellular pathway^63, 64^. It will be interesting to determine whether RBPs are similarly modified between serotypes or whether effects on RNA processing are achieved through different substrates. There is a precedent for post-translational modification regulating hnRNP-C affinity for RNA, with conjugation of the ubiquitin-like protein SUMO decreasing the affinity of hnRNP-C for RNA^65^. Ubiquitin and related proteins have emerging roles in regulating splicing by altering the properties and dynamics of spliceosomal complexes through altered protein-protein interactions^66^. It is likely that RBPs such as RALY and hnRNP-C are also functionally regulated through ubiquitination by cellular ubiquitin ligases. Correlating changes to host splicing induced as a result of the impact of ubiquitination during Ad infection may provide insights into host pathways that are altered by ubiquitination of these RBPs. In addition to the ubiquitin-mediated changes in interaction of hnRNP-C and RALY with viral late RNA, we also observed global changes in the RNA-binding of hnRNP-C during infection, independent of the Ad ubiquitin ligase complex (**Fig. 5**). Understanding how Ad induces the reduction of RNA-binding by the hnRNP-C RRM and the associated changes in binding motif, may provide novel insights into regulation of RBP function.

Manipulation of the host ubiquitin machinery during virus infection has traditionally been studied in the context of proteasomal degradation and there are very few known examples of viruses directing ubiquitin towards cellular substrates that are not subsequently degraded^1–3^. This has been true for the Cullin5 ligase redirected by Ad E1B55K/E4orf6 which was previously shown to induce degradation of proteins involved in the cellular DNA damage response and apoptosis^8, 19–21, 23^. Our observation that the majority of potential cellular substrates of the E1B55K/E4orf6 complex appear to be ubiquitinated without significant decrease in abundance suggests that a major aspect of the activity of the viral assembled ligase is non-degradative ubiquitination. This finding highlights the need to combine ubiquitinome analysis together with whole cell proteome quantification when identifying outcomes of ubiquitination. Future studies of other viral ligases should include this type of analysis of non-degradative ubiquitination in order to ensure that all aspects of viral manipulation by ubiquitin are identified. We propose that ubiquitination without the need for proteasome-mediated degradation provides increased flexibility and more rapid approaches to counter host responses and redirect cellular processes. Viral redirection of ubiquitination may present particularly good model systems to study how ubiquitin ligases in general can facilitate both degradative and non-degradative ubiquitination of distinct substrates. Given the increasing appreciation that cellular ubiquitin ligases (such as Cullin ligases) can facilitate the formation of multiple different types of ubiquitin chains^67–69^, viral infections provide systems to decipher the rules that govern outcomes of ubiquitination.

In addition to its contributions to fundamental knowledge of cellular and molecular biology, Ad has also been developed as a vector for gene delivery and oncolytic cancer treatment. Mutant viruses that lack E1B55K have been shown to replicate conditionally in cancer cells, with selectivity that was initially suggested to be based on p53 inactivation but is more likely due to preferential viral late mRNA export^70–72^. Since many cancers have altered RNA processing, the Ad ΔE1B used for oncolytic therapies may be complemented by defects in substrates of the E1B55K/E4orf6 complex. Our work suggests that alterations in these substrates, such as the RBPs RALY and hnRNP-C, may make tumor cells more susceptible to ΔE1B-based oncolytic viruses.

## Materials and Methods

### Cell culture

All cell lines were obtained from the American Type Culture Collection (ATCC) and cultured at 37°C and 5% CO_2_. HeLa (Cat#: ATCC CCL-2), HEK293 (Cat#: ATCC CRL-1573), and U2OS cells (Cat#: ATCC HTB-96) were grown in DMEM (Corning, Cat#: 10-013-CV) supplemented with 10% v/v fetal bovine serum (FBS) (VWR, Cat#: 89510-186) and 1% v/v Pen/Strep (100 U/ml of penicillin, 100 μg/ml of streptomycin, Gibco, Cat#: 15140-122). A549 cells (Cat#: ATCC CCL-185) were maintained in Ham’s F-12K medium (Gibco, Cat#: 21127-022) supplemented with 10% v/v FBS and 1% v/v Pen/Strep. Primary like HBEC3-KT (Cat#: ATCC CRL-4051) were grown in Airway Epithelial Cell Basal Medium (Cat#: ATCC PCS-300-030) supplemented with Bronchial Epithelial Cell Growth Kit (Cat#: ATCC PCS-300-040) and 1% v/v Pen/Strep. The RALY-Flag inducible cell line was generated using a HeLa acceptor cell line kindly provided by E. Makeyev^73^ and used as previously reported. RALY-Flag was cloned from the pcDNA3.1 plasmid described below and inserted into the inducible plasmid cassette using restriction enzymes BsrGI and AgeI. Sequence confirmed clones were transfected into the HeLa acceptor cells along with plasmid encoding the Cre recombinase. Clones were selected by puromycin (1 µg/mL) and induced with doxycycline (0.5 µg/mL) to express RALY-Flag. Protein expression was verified by immunoblot. All cell lines tested negative for mycoplasma using the LookOut Mycoplasma PCR Detection Kit (Sigma-Aldrich).

### Viruses and infection

Ad5 wild-type (WT) was purchased from ATCC. The Ad5 E1B55K-deletion mutant dl110 has been described previously^10^ and was a gift from G. Ketner. The E1 deletion mutant recombinant adenovirus vectors expressing E1B55K (rAd E1B55K)^35^ and E4orf6 (rAd E4orf6)^19^ were obtained from P. Branton. All viruses were propagated on HEK293 cells, purified using two sequential rounds of ultracentrifugation in CsCl gradient and stored in 40% v/v glycerol at −20°C. Viral titers were determined by plaque assay on HEK293 cells for all but rAd E4orf6. For this virus we assumed a plaque forming unit-to-particle ratio of 1:50. All infections were carried out using a multiplicity of infection (MOI) of 10 and harvested at indicated hours post infection (hpi). Infections were performed on monolayers of cells by dilution of the virus in respective low serum growth medium. After 2 h at 37°C additional full serum growth medium was added. For plaque assays, the virus infection media was removed after 2 h and cells were washed 1x in PBS before addition of full serum growth medium.

### Plasmids, siRNA and transfection

Full-length RALY with a carboxyl-terminal Flag-tag (cDNA obtained from Dharmacon, Cat#: MHS6278-202857995) and hnRNP-C isoforms 1 and 2 with a carboxyl-terminal Flag-tag (cDNA containing plasmids were a gift from K. Lynch) and RFP were cloned into the pcDNA3.1 vector using the BamHI and XbaI restriction sites. The pRK5 vector encoding Ad5 E4orf6 was generated by subcloning from purified Ad5 DNA as previously described^74^. The expression vector for HA-tagged tetra-ubiquitin as previously described^75^ was a gift from R. Greenberg DNA transfections were performed using the standard protocol for Lipofectamine2000 (Invitrogen).

The following siRNAs were obtained from Dharmacon: non-targeting control (Cat#: D-001206-13-05), RALY (Cat#: M-012392-00-0005) and hnRNP-C (Cat#: M-011869-01-0005; Cat#: L-011869-03-0005 only used for hnRNP-C single knockdown in supplementary Fig. 7). siRNA transfections were performed using the standard protocol for Lipofectamine RNAiMAX (Invitrogen).

### Antibodies and inhibitors

The following primary antibodies for viral proteins were obtained: Adenovirus late protein antibody staining Hexon, Penton and Fiber (gift from J. Wilson^76^, species: rabbit, WB 1:10,000), Protein VII (gift from H. Wodrich^77^, Clone: Chimera 2-14, WB 1:200), DBP (gift from A. Levine^78^, Clone: B6-8, WB 1:1000, IF 1:400), E1B55K (gift from A. Levine^79^, Clone: 58K2A6, WB 1:500) and E4orf6 (gift from D. Ornelles^80^, Clone: RSA#3, WB 1:500). The following primary antibodies were used for cellular proteins: MRE11 (Novus Biologicals, Catalog#: NB100-142, WB 1:1000), BLM (Abcam, Catalog#: ab476, WB 1:1000), Cullin5 (Bethyl Laboratories, Catalog#: A302-173A, WB 1:200), Actin (Sigma-Aldrich, Catalog#: A5441-100UL, WB 1:5000), RALY (Bethyl Laboratories, Catalog#: A302-070A, WB 1:1000; Bethyl Laboratories, Catalog#: A302-069A, IF 1:500, IP 5 μl = 5 μg), hnRNP-C (Santa Cruz Biotechnology, Catalog#: sc-32308, WB 1:1000, IF 1:1000, IP 25 μl=5 μg), Tubulin (Santa Cruz Biotechnology, Catalog#: sc-69969, WB 1:1000), Flag (Sigma-Aldrich, Catalog#: F7425-.2MG, WB 1:1000; Sigma-Aldrich, Catalog#: F3165-1MG, IP 5 μg), Ubiquitin (Santa Cruz, Catalog#: sc-9133, IP 10 μl=2 μg; Abcam, Catalog#: ab7780, IP 5 μl), NBS1 (Novus Biologicals, Catalog#: NB100-143, WB 1:1000), RAD50 (GeneTex, Catalog#: GTX70228, WB 1:1000) and USP7 (Bethyl Laboratories, Catalog#: A300-033A, IF 1:500).

Horseradish peroxidase-conjugated (HRP) secondary antibodies for immunoblot were purchased from Jackson Laboratories. Anti-mouse IgG conjugated to HRP for immunoblot of immunoprecipitation samples (used in Fig. 3b) was purchased from Abcam (Cat#: ab131368). Fluorophore-conjugated secondaries for immunofluorescence were purchased from Life Technologies.

Cycloheximide (CHX) was purchased from Calbiochem (Cat#: 293764), dissolved in DMSO to a stock concentration of 25 mM and used at a final concentration of 25 μM. NEDDylation inhibitor MLN4924 was purchased from Sigma-Aldrich (Cat#: 505477), dissolved in DMSO to a stock concentration of 1 mM and used at a final concentration of 3 μM. Proteasome inhibitor MG132 was purchased from Sigma-Aldrich (Cat#: 474791) at a concentration of 10 mM in DMSO and used at a final concentration of 20 μM.

### Immunoblotting

Protein samples were prepared using lithium dodecyl sulfate (LDS) loading buffer (NuPage) supplemented with 25 mM dithiothreitol (DTT) and boiled at 95°C for 10 min. Equal amounts of protein lysate were separated by SDS-PAGE and transferred onto a nitrocellulose membrane (Millipore) at 30 V for at least 60 min (overnight for ubiquitination assays). Membranes were stained with Ponceau to confirm equal loading and blocked in 5% w/v milk in TBST supplemented with 0.05% w/v sodium azide. Membranes were incubated with primary antibodies overnight, washed for 30 min in TBST, incubated with HRP-conjugated secondary for 1 h and washed again for 30 min in TBST. Proteins were visualized with Pierce ECL Western Blotting Substrate (Thermo Scientific) and detected using a Syngene G-Box. Images were processed and assembled in Adobe CS6. Immunoblots were quantified by pixel densitometry using the Syngene GeneTools software.

### Immunofluorescence

HeLa cells were grown on coverslips in 24-well plates, infected with indicated viruses and fixed at 24 hpi in 4% w/v paraformaldehyde in PBS for 10 mins. Cells were permeabilized with 0.5% v/v Triton-X in PBS for 10 mins. The samples were blocked in 3% w/v BSA in PBS (+ 0.05% w/v sodium azide) for 30 mins, incubated with primary antibodies in 3% w/v BSA in PBS (+ 0.05% w/v sodium azide) for 1 h, followed by secondary antibodies and 4,6-diamidino-2-phenylindole (DAPI) for 2 h. Secondary antibodies used were Alexa Fluor α-rabbit 488 and α-mouse 555. Coverslips were mounted onto glass slides using ProLong Gold Antifade Reagent (Cell Signaling Technologies). Immunofluorescence was visualized using a Zeiss LSM 710 Confocal microscope (Cell and Developmental Microscopy Core at UPenn) and ZEN 2011 software. Images were processed in ImageJ and assembled in Adobe CS6.

### RNA Fluorescence *in situ* hybridization

RNA FISH was performed following previously established protocols^81^, with the following modifications. Thirty-two singly labeled DNA oligonucleotides targeting the Fiber open reading frame were designed using the Stellaris smFISH probe designer and ordered with a 3’ mdC-TEG-Amino label from LGC Biosearch. Fiber FISH probes were pooled and labeled with ATTO 647N NHS-Ester (ATTO-TEC, Cat#: AD 647N-31), isopropanol precipitated and purified by HPLC as previously described^81^. GAPDH probes labeled with Cy3 were used as a counterstain to demarcate cytoplasmic boundaries and were a kind gift from Sydney Schaffer, University of Pennsylvania^82^. All probe sequences can be found in **Supplementary Table 7**. HeLa cells were grown on coverslips, harvested, fixed, and permeabilized as described for conventional immunofluorescence above. After permeabilization, cells on coverslips were equilibrated in Wash Buffer (2X SSC, 10% formamide) before being inverted over 30 μl Hybridization Buffer (2X SSC, 10% formamide, 10% dextran sulphate) containing 500 nM Fiber and GAPDH FISH probes and incubated at 37°C in a humidified chamber overnight. The following day coverslips were washed twice with Wash Buffer for 30 minutes at 37°C with DAPI added to the second wash, briefly washed three times at room temperature with 2X SSC, and then affixed to glass slides using clear nail polish. Images were acquired on a Zeiss LSM 710 microscope with ten z-stacks of 0.7 μm each in the z-direction. Images were deconvoluted by maximum intensity projection in the z-direction using ImageJ. Fiber RNA localization was scored as described in **Supplementary Figure 1** over 41-160 individual cells. Representative images were further processed in ImageJ and assembled in Adobe CS6.

### RNA isolation and RT-qPCR

Total RNA was isolated from infected cells at the indicated timepoints using the RNeasy Micro Kit (Qiagen). Complementary DNA (cDNA) was synthesized using 1 μg of input RNA and the High Capacity RNA-to-cDNA Kit (Thermo Fisher). Quantitative PCR was performed by standard protocol using diluted cDNA, primers for different viral and cellular transcripts (see **Supplementary Table 7** for complete list of primers) and SYBR Green (Thermo Scientific) using the QuantStudio 7 Flex Real-Time PCR System (Thermo Scientific). The relative values for each transcript were normalized to a control RNA (actin or HPRT).

### RNA Transcription and Stability Profiling

To assess relative RNA transcription rate and RNA half-life, cells were treated with 200 μM 4-thiouridine (4sU; Sigma T4509) for exactly 30 min. Infection was stopped and RNA harvested using 1 ml TRIzol (Thermo Fisher Scientific), following manufacturer’s instructions. A fraction of the total RNA was reserved as input, and the remaining 4sU-labeled nascent RNA was biotinylated using MTSEA-Biotin-XX (Biotium; 90066) as previously described^83, 84^. Nascent RNA was separated from unlabeled RNA using MyOne C1 Streptavidin Dynabeads (Thermo Fisher Scientific; 65-001), biotin was removed from nascent RNA using 100 mM dithiothreitol (DTT), and RNA was isopropanol precipitated. Total RNA (1 μg) and an equivalent volume of nascent RNA were converted to cDNA and qPCR was performed as described above. Relative transcription rates were determined by the ΔΔCt method to compare nascent transcript levels between control and siRNA treated cells normalized to nascent GAPDH RNA. RNA half-life was determined using the previously described formula *t*_1/2_ = -*t* × [ln(2)/*DR*] where *t* is the 4sU labeling time (0.5 h) and *DR* is the decay rate defined as Nascent/Total RNA^85^. Half-lives were normalized to the half-life of GAPDH set at 8 h as previously determined^86^.

### RNA decay measurement using Actinomycin D

To determine the decay of viral mRNA species, HeLa cells infected with either Ad5 WT or ΔE1B were treated with 10 μM Actinomycin D (Cayman Chemical, Cat#: 11421) at 24 hpi. RNA harvested using RLT buffer (from Qiagen RNA isolation kit) at 0, 1, 2, 4, 6, and 8 hours after treatment. RNA levels were quantified using RT-qPCR and normalized to 0 hours of Actinomycin D to determine RNA decay.

### Viral genome accumulation by qPCR

Infected cells were harvested by trypsinization at 4 and 24 hpi and total DNA was isolated using the PureLink Genomic DNA kit (Invitrogen). qPCR was performed using primers for the Ad5 DBP and cellular tubulin (see **Supplementary Table 7** for primers). Values for DBP were normalized internally to tubulin and to the 4 hpi timepoint to control for any variations in virus input. qPCR was performed using the standard protocol for SYBR Green and analyzed with the QuantStudio 7 Flex Real-Time PCR System.

### Plaque assay

Infected cells seeded in 12-well plates were harvested by scraping at the indicated timepoints and lysed by three cycles of freeze-thawing. Cell debris was removed from lysates by centrifugation at max speed (21,130 g), 4°C, 5 min. Lysates were diluted serially in DMEM supplemented with 2% v/v FBS and 1% v/v Pen/Strep to infect HEK293 cells seeded in 12-well plates. After incubation for 2 h at 37°C, the infection media was removed, and cells were overlaid with DMEM containing 0.45% w/v SeaPlaque agarose (Lonza) in addition to 2% v/v FBS and 1% v/v Pen/Strep. Plaques were stained using 1% w/v crystal violet in 50% v/v ethanol between 6 to 7 days post-infection.

### Immunoprecipitation

Approximately 2×10^7^ cells were harvested, washed, pelleted and flash frozen for each immunoprecipitation. For IP of hnRNP-C and RALY 50 μl of Protein G Dynabeads (Thermo Fisher) per sample were washed 3x in IP buffer (50 mM HEPES pH 7.4, 150 mM KCl, 2 mM EDTA, 0.5% v/v NP-40, 0.5 mM DTT, 1x cOmplete Protease Inhibitor Cocktail (Roche)) and incubated with 5 μg of antibody (α-hnRNP-C or α-RALY) rotating at 4°C for 2h. Cell pellets were resuspended in 1 ml IP buffer and incubated for 1 h on ice. Samples were sonicated with a Diagenode Biorupter on low setting for 30 s on and 30 s off for ten rounds at 4°C and spun at max speed (21,130 g) for 10 min at 4°C. 300 μl of sample were added to washed beads and incubated rotating at 4°C for 2h. Beads were washed 4x in IP wash (same as above but with only 0.05% v/v NP-40). Samples were eluted in 50 μl 1x LDS sample buffer with 25 mM DTT by boiling for 10 min at 95°C and further processed for analysis by immunoblot. The following changes were made to the protocol for IP of E1B55K: IP buffer contained 50 mM Tris-HCl pH 7.4, 0.1% v/v Triton X-100, 150 mM NaCl, 50 mM NaF, 1 mM Na_3_VO_4_, 1x cOmplete Protease Inhibitor Cocktail.

### Denaturing *in vivo* ubiquitination assay

Approximately 1×10^7^ cells were washed, pelleted and stored at −80°C for each immunoprecipitation. For HEK293 cells, the pellets were thawed on ice and resuspended in 100 μl of Lysis buffer (1% w/v SDS, 5 mM EDTA, 10 mM DTT, 1x cOmplete Protease Inhibitor Cocktail) with 1 μl Benzonase (Sigma-Aldrich) by vortexing. Samples were incubated on ice for 5 min and then further denatured by heating to 95°C for 5 min. 900 μl of Wash buffer (10 mM Tris-HCl pH 7.4, 1 mM EDTA, 1 mM EGTA, 150 mM NaCl, 1% v/v Triton X-100, 0.2 mM Na_3_VO_4_, 1x cOmplete Protease Inhibitor Cocktail), passed 10 times through a 23G syringe and spun at max speed (21,130 g) for 5 min at 4°C. A minimum of 800 μl of sample was added to 50 μl washed Pierce Anti-HA Magnetic beads (Thermo Fisher). Sample was incubated with beads rotating for 1 h at 4°C, washed 3x in Wash buffer and eluted in 1x LDS sample buffer with 25 mM DTT for further processing by immunoblot.

The following changes were made to the protocol for HeLa cells: the Lysis buffer contained 1% w/v SDS in PBS, Tris buffered saline with Tween-20 was used as wash buffer, Protein G Dynabeads incubated for 1 h with a mix of both α-ubiquitin antibodies listed above were used for the IP.

### De-ubiquitination assay

Approximately 1×10^7^ HEK293 cells were washed, pelleted and stored at −80°C for each immunoprecipitation. The pellets were resuspended in 1 ml IP buffer B (20 mM HEPES-KOH pH 7.4, 110 mM potassium acetate, 2 mM MgCl_2_, 0.1% v/v Tween-20, 0.1% v/v Triton X-100, 150 mM NaCl, 1 mM DTT, 0.1 mM PTSF) containing 1x cOmplete Protease Inhibitor Cocktail, 20 μM PR-619 (LifeSensors, Cat#: SI9619-5X5MG), 5 mM 1,10-phenanthroline (LifeSensors, Cat#: SI9649), and 1 μl/ml Benzonase (Sigma-Alrich). Samples were incubated on ice for 30 min, sonicated with a Diagenode Biorupter on low setting for 30 s on and 30 s off for five rounds at 4°C and spun at max speed (21,130 g) for 5 min at 4°C. 925 μl of sample was added to 100 μl washed Pierce Anti-HA Magnetic beads (Thermo Fisher). Sample was incubated with beads rotating for 2 h at 4°C, washed 3x in IP buffer B, resuspended in 100 μl of IP buffer B and split into three 30 μl aliquots. 1 μl of 20 mM PR-619 was added to sample 1 (untreated), 1 μl of USP2 (LifeSensors, Cat#: DB501) was added to sample 2 (DUB^PAN^) and 2 μl of OTUB1 (LifeSensors, Cat#: DB201) was added to sample 3 (DUB^K48^). Samples were incubated at 30°C for a minimum of 1 h. Samples were eluted by addition of 10 μl 4x LDS sample buffer with 100 mM DTT and boiling at 95°C for 10 min for further processing by immunoblotting.

### CLIP-qPCR

The CLIP protocol was adapted from existing protocols^58^. In short, approximately 2×10^7^ cells were crosslinked on ice with 0.8 J/cm^2^ UV 254 nm in a UV Stratalinker 2400 (Stratagene), washed in PBS with 2 mM EDTA and 0.2 mM PMSF, flash frozen in liquid nitrogen and stored at −80°C. 50 μl of Protein G Dynabeads per sample were washed 3x in iCLIP lysis buffer A (50 mM Tris-HCl pH 7.4, 100 mM NaCl, 0.2% v/v NP-40, 0.1% w/v SDS, 0.5% w/v Sodium deoxycholate, 1x cOmplete Protease Inhibitor Cocktail), resuspended in 100 μl iCLIP lysis buffer A and incubated with 5 μg of α-hnRNP-C antibody, 5 μg of α-Flag antibody (mouse), or 5 μl of Normal Mouse Serum Control (Thermo Fisher) rotating 1 h at 4°C. Cell pellets were resuspended in 1 ml of iCLIP lysis buffer B (same as buffer A but with 1% v/v NP-40 and 11 μl of Murine RNase inhibitor (NEB) per 1 ml) and incubated on ice for 15 min. Samples were sonicated with a Diagenode Biorupter on low setting for 30 s on and 30 s off for five rounds at 4°C. 2 μl of TURBO DNase (Thermo Fisher) were added and samples incubated at 37°C for 6 min. Lysates were cleared by centrifugation at max speed (21,130 g) for 15 min at 4°C and supernatants transferred to a new tube. 300 μl of lysate were added to washed beads and incubated rotating at 4°C for 2 h. Beads were washed 2x in High Salt buffer (50 mM Tris-HCl pH 7.4, 1 M NaCl, 1 mM EDTA, 0.2% v/v NP-40, 0.1% w/v SDS, 0.5% w/v Sodium deoxycholate), 2x in Wash buffer (20 mM Tris-HCl pH 7.4, 10 mM MgCl_2_, 0.2% v/v Tween-20) and 2x in Proteinase K buffer (100mM Tris-HCl pH 7.4, 50 mM NaCl, 10 mM EDTA, 0.2% w/v SDS). Beads were resuspended in 50 μl Proteinase K buffer and 10 μl removed and processed for immunoblot analysis. 10 μl of Proteinase K (NEB) and 2 μl Murine RNase Inhibitor were added to the remaining beads or to 30 μl of input (10%) and incubated at 50°C for 1 h. The RNA was extracted using a standard protocol for TRIzol (Thermo Fisher) and further processed for RT-qPCR.

### seCLIP-Seq

#### Sample preparation

The CLIP protocol was adapted from existing protocols^87^. In short, approximately 2×10^7^ HeLa cells were crosslinked on ice with 0.8 J/cm^2^ UV 254 nm in a UV Stratalinker 2400 (Stratagene), washed in PBS with 2 mM EDTA and 0.2 mM PMSF, flash frozen in liquid nitrogen and stored at −80°C. Protein G Dynabeads (100 μl per sample) were washed 3x in iCLIP lysis buffer A (see CLIP-qPCR), resuspended in 100 μl iCLIP lysis buffer A and incubated with 10 μg of α-hnRNP-C antibody rotating 1 h at RT. Cell pellets were resuspended in 1 ml of iCLIP lysis buffer B and incubated on ice for 15 min. Samples were sonicated with a Diagenode Biorupter on low setting for 30 s on and 30 s off for five rounds at 4°C. Samples were incubated with 2 μl of TURBO DNase (Thermo Fisher) and 10 μl of 1:10 diluted RNase I (Thermo Fisher) in Thermomixer at 1200 rpm at 37°C for 5 min, samples placed on ice and 22 μl SUPERase·In RNase Inhibitor was added. Cleared lysate (500 μl) was added to washed beads and incubated rotating at 4°C for 2 h. Beads were washed 2x in High Salt buffer, 2x in Wash buffer and 2x FastAP buffer. FastAP master mix (100 μl) and FastAP enzyme (8 μl) was added and samples were incubated with a Thermomixer at 1200 rpm at 37°C for 15 min. T4 PNK enzyme (7 μl) and 300 PNK master mix were added and samples were incubated with a Thermomixer at 1200 rpm at 37 °C for 20 min. Beads were washed and resuspended in Ligase buffer with 2.5 μl RNA Ligase high conc., 2.5 μl of RNA adapters (3SR_RNA), and incubated at RT for 75 min. Beads were washed and a fraction saved for immunoblotting. For the remaining fraction, beads were resuspended in lysis buffer with DTT, eluted by incubation in Thermomixer, 1200 rpm, 70 °C, run on SDS-PAGE and transferred onto Nitrocellulose o/n, 30V. Lanes for the RBP band (plus 75 kDa) and size-matched input were cut from the membranes RNA was eluted with 20 μl of Proteinase K (Thermo Fisher) in a Thermomixer at 1200 rpm at 50 °C for 1 h. RNA was extracted with acid phenol/chloroform/isoamyl alcohol (pH 4.5), and concentrated using RNA Clean and Concentrator (Zymo). Size-matched inputs were ligated to 3SR_RNA. All RNA samples were reverse transcribed with 0.9 μl AffinityScript Enzyme at 55 °C for 45 min with RT primer SR_RTv2. RNA and excess primers were removed with 3.5 μl ExoSAP-IT 1 M NaOH. cDNA was purified using 10 μl MyONE Silane beads and ligated to 5’ linker SR_DNA o/n at RT. After clean up, cDNA was quantified by qPCR using NEBNext universal and index primers (NEB E7335S). Libraries were indexed using NEBNext High-Fidelity PCR Master Mix (NEB M0541S) for 11 cycles (size-matched input) or 15 cycles (hnRNP-C IP). Libraries were size selected by 1.0x AmpureXP beads (Beckman Coulter A63880), quantified by QuBit HS DNA and Bioanalyzer High Sensitivity DNA assay, and pooled for sequencing.

#### Data analysis

Preprocessing involved adapter cutting using cutadapt (v. 1.18)^88^ and extracting the UMIs using umi_tools (v 1.0.0)^89^. Alignment was achieved using GSNAP (v 2019-09-12)^90^. Reads were aligned to the human and adenovirus 5 genome simultaneously. After the alignment we used umi_tools to deduplicate reads based on the UMIs, which was followed by removing all non-unique reads. We then used clipper (v0.1.4)^91^ to find significant enriched IP peaks over the input on the human genome. To identify enriched peaks on the virus genome we employed a sliding window approach, by counting fragments overlapping 10bp wide windows along both the forward and reverse strand on the genome. If two consecutive windows were significantly enriched over input, they were merged into one peak. Motif analysis was conducted using the Homer suite^92^.

### Analyzing protein complexes by crosslinking

Cells were crosslinked using disuccinimidyl suberate (DSS, Thermo Fisher) dissolved to 100 mM in DMSO and further diluted to 0.1 mM, 0.3 mM and 1 mM in PBS. Cells seeded as a monolayer in 6-well plates were washed once with PBS, overlaid with 500 μl with PBS or the different DSS dilutions and incubated at room temperature for 30 min. The reaction was quenched by addition of 500 μl of 20 mM Tris-HCl pH 7.4, washed twice with PBS and further processed for immunoblot analysis.

### Di-glycine remnant profiling by mass spectrometry

#### Cell lysis and initial desalting

Approximately 10 mg of input was generated from 5×15 cm plates of HeLa cells transduced with rAd E1B55K and rAd E4orf6 constructs for 0 h (mock), 6 h, 8 h, and 10 h. Each timepoint was produced in biological triplicate. Cell were harvested with 0.25% Trypsin (Gibco), washed 1x in PBS, flash frozen in liquid nitrogen and stored at −80°C. Pellets were thawed, resuspended in 1 ml of lysis buffer (6 M urea, 2 M thiourea, in 50 mM ammonium bicarbonate pH 8.0) with 1x Halt Protease Cocktail inhibitor solution, and incubated for ∼5 min on ice. Samples were then diluted 10-fold in 50 mM ammonium bicarbonate, reduced with 10 mM DTT, alkylated with 20 mM iodoacetamide, and digested with trypsin protease overnight. Digestion was quenched by acidification to pH 2 with trifluoroacetic acid (TFA) and samples were desalted over Waters tC18 SepPak cartridges (Cat#: 036805). A 10% aliquot was set aside for global proteomic analysis and all samples were dried to completion.

#### Di-glycine (K-ε-GG) enrichment, fractionation, and desalting

A Cell Signaling PTMScan ubiquitin remnant motif kit (Cat#: 5562) was used to enrich for peptides that had been ubiquitinated. Aliquoted beads were cross-linked for 30 minutes in 100 mM sodium borate and 20 mM dimethyl pimelimidate (Thermo Scientific), following the protocol outlined by Udeshi *et. al*.^28^. Tryptic peptides were resuspended in IAP buffer (50 mM MOPS, pH 7.2, 10 mM sodium phosphate, 50 mM NaCl) and immunoprecipitated with the provided antibody for 2 h at 4°C. Samples were eluted in LC-MS grade water (Thermo Fisher) with 0.15% v/v TFA and separated into either 3 high-pH fractions (enriched ubiquitinated peptides) or 7 high-pH fractions (global proteome) over C18 columns (The Nest Group, MicroSpin column C18 silica, part#: SEM SS18V, lot#: 091317). Fractionated samples were desalted a final time over Oligo R3 reverse-phase resin (Thermo Scientific, Cat#:1-1339-03).

#### Data acquisition and search parameters

All solvents used in analysis of MS samples were LC-MS grade. Samples were analyzed with an Easy-nLC system (Thermo Fisher) running 0.1% v/v formic acid (Buffer A) and 80% v/v acetonitrile with 0.1% v/v formic acid (Buffer B), coupled to an Orbitrap Fusion Tribrid mass spectrometer. Peptides were separated using a 75 µm i.d. silica capillary column packed in-house with Repro-Sil Pur C18-AQ 3 µm resin and eluted with a gradient of 3-38% Buffer B over 85 minutes. Full MS scans from 300-1500 m/z were analyzed in the Orbitrap at 120,000 FWHM resolution and 5×10^5^ AGC target value, for 50 ms maximum injection time. Ions were selected for MS2 analysis with an isolation window of 2 m/z, for a maximum injection time of 50 ms, to a target AGC of 5×10^4^.

MS raw files were analyzed by MaxQuant software version 1.6.0.16, and MS2 spectra were searched against a target + reverse database with the Andromeda search engine using the Human UniProt FASTA database [9606] (reviewed, canonical entries; downloaded November 2017) and adenovirus serotype 5 UniProt FASTA database (reviewed, canonical entries; downloaded February 2018). The search included variable modifications of methionine oxidation, N-terminal acetylation, and GlyGly on lysine residues, with a fixed modification of carbamidomethyl cysteine. For global proteome samples, iBAQ quantification was performed on unique+razor peptides using unmodified, oxidized methionine, and N-terminally acetylated forms. Trypsin cleavage was specified with up to 2 missed cleavages allowed. Match between runs was enabled, but restricted to matches within a single biological replicate by separating replicates into independent searches. Match between runs parameters included a retention time alignment window of 20 min and a match time window of 0.7 min. False discovery rate (FDR) was set to 0.01.

### Proteomics and bioinformatics analysis

#### Data normalization and filtering

MaxQuant output was filtered to remove identified contaminant and reverse proteins. MaxQuant “Intensity” and “iBAQ”^93^ label free quantification values were used to measure abundances for the K-ε-GG and WCP data, respectively. Abundances were transformed to log2 values, with unidentified values assigned as “NA”. K-ε-GG and WCP data were normalized separately. Data were normalized by subtracting the sample medians from log2 transformed abundances within replicates. Both the KεGG and the WCP datasets were filtered at each timepoint to require quantification in at least 2 of 3 replicates to be included in the calculations of fold change, z-score, or for hypothesis testing. Data that contained less than 2 replicate quantifications in each timepoint was removed entirely from the analysis. Peptides or proteins are considered uniquely identified in one timepoint compared to another if there were at least 2 replicate quantifications for one timepoint and 0 replicate quantifications for the compared timepoint.

#### Fold change, p-value, and z-score calculations

The fold change across timepoints was calculated by comparing the log2 transformed, normalized peptide or protein abundances for compared timepoints. Fold changes were calculated both on a per-replicate basis and by comparing averaged abundances across timepoints. Hypothesis testing was performed using unpaired, two-tailed Students t-tests, when comparing log2 transformed, normalized replicate abundances across timepoints. Hypothesis testing using one-sided t-tests, with null hypothesis of fold change equal 0, was implemented when evaluating log2 fold changes. Multiple testing correction was not performed. Peptide ubiquitin intensity Z-scores were used to measure relative ubiquitin abundances for a peptide at the respective timepoint. Z-scores were calculated by averaging the peptide intensities for each replicate identification within the timepoint and comparing to the mean and standard deviation of averaged values within that timepoint.

#### Protein ubiquitin abundance calculation

The di-glycine technique quantifies peptide-based abundance of the K-ε-GG modification. In order to quantify protein-based K-ε-GG abundance changes, we implemented a calculation to combine the peptide-based fold changes for cases in which multiple K-ε-GG peptides comprise a modified protein. If a single K-ε-GG peptide was identified for a modified protein, that K-ε-GG peptide abundance fold change represented the protein-based K-ε-GG fold change. For cases in which multiple K-ε-GG peptides were quantified for a single protein, the fold changes of each peptide were weighted by the abundance of that peptide and the weighted fold changes were averaged to calculate the protein-based K-ε-GG fold change. In cases for which a peptide was uniquely identified in the mock or 10 h transduction timepoint, a log2 fold change of plus or minus 7, respectively, representing the largest fold changes identified in the dataset, was assigned to this peptide. The K-ε-GG abundance log2 fold changes, for each identified replicate, were normalized by the total protein abundance log2 fold change of the corresponding replicate of the same protein in the corresponding whole cell proteome. The replicate-based normalized log2 fold changes were averaged and hypothesis testing was performed for the log2 fold changes using onesided t-tests. The normalization of the K-ε-GG fold change by total protein fold change was performed to identify differentially increased or decreased ubiquitination, beyond what would be expected if modification abundance was driven solely by changes in total protein abundance.

#### K-ε-GG and whole cell proteome comparison

The protein-based K-ε-GG and corresponding whole cell proteome data were compared to identify proteins that exhibited an increase in K-ε-GG abundance and to predict the effect of ubiquitination on total protein abundance. Proteins that exhibited a protein-based, normalized K-ε-GG log2 fold change > 1 were classified as being increased in ubiquitination in response to E1B55K/E4orf6 expression. Proteins that exhibited whole cell proteome log2 fold change greater than the mean fold change +/- 1 standard deviation, or which were uniquely identified in the 0 or 10 hour timepoint, were classified as increased or decreased in total protein abundance. Proteins for which total protein expression did not deviate more than +/- 1 standard deviation from the mean fold change were classified as unchanged in protein abundance in response to E1B55K/E4orf6 expression. Proteins that were ubiquitinated and decreased in total protein abundance were predicted to be potential substrates of E1B55K/E4orf6 ubiquitin-mediated degradation. Proteins that were ubiquitinated and unchanged in total protein abundance were predicted to be non-degraded substrates of E1B55K/E4orf6.

#### Gene ontology and protein-protein interaction network analysis

The proteins that exhibited increased protein-based ubiquitination were analyzed using the ReactomeFI plug-in (6.1.0)^36^ within the Cytoscape network visualization software (3.4.0)^94^. The protein-protein interaction network was generated using the Gene Set analysis within the “2016” ReactomeFI network version with “linker genes” included. The network was clustered using the in-built ReactomeFI clustering algorithm. Gene ontology “Molecular Function”, “Biological Processes” and Reactome Pathway analysis was performed within the ReactomeFI application for the entire network as well as for each clustered module. Network node attributes included size, which corresponded to degree of increased ubiquitination, and color, which corresponded to total protein increase or decrease. Network edges were set to non-directed, solid lines for all types of Reactome protein-protein interactions.

### Targeted hnRNP-C RBR-ID

#### Cell growth

Heavy and light media were prepared by supplementing SILAC DMEM (Thermo #88364) with 800 µM of Lysine (Sigma #L8662-25G) and 400 µM Arginine (Sigma #A8094-25G) for light or K8 (Silantes #211604102) and R10 isotopes (Silantes #201604102) for heavy, and 120 mg/L Proline (Sigma #P0380-100G). Media was then filtered and adjusted to 10% dialyzed FBS (HyClone #SH30079.03) and 1% penicillin-streptomycin. Heavy isotope labeling in HeLa cells was confirmed by mass spectrometry. Cells were either mock-treated or infected with Ad5 WT Ad5 or ΔE1B at an MOI of 10. At 20 hpi, media was exchanged and heavy-labeled cells were pulsed with 500 µM 4sU, with light-labeled cells serving as non-treated controls. At 24 hpi, all samples were washed with cold PBS and crosslinked at 1.0 J/cm^2^ with 310 nm UV-B. Cells were then harvested, heavy/light pairs were combined 1:1, and aliquoted for further analysis.

#### hnRNP-C IP

Approximately 1×10^7^ pooled heavy and light HeLa cells were used for one hnRNP-C IP. 50 μl of Protein G Dynabeads (Thermo Fisher) per sample were washed 3x in IP buffer (20 mM HEPES-KOH pH 7.4, 110 mM potassium acetate, 2 mM MgCl_2_, 0.1% Tween-20, 0.1% Triton, 150 mM NaCl, 1 mM DTT, 0.1 mM PMSF, 1x cOmplete Protease Inhibitor Cocktail (Roche)) and incubated with 5 μg of α-hnRNP-C antibody rotating at RT for 1 h. Cell pellets were resuspended in 500 μl IP buffer, after 10 min 1.5 µl benzonase (Sigma-Aldrich, Cat#: E1014) were added and the sample was incubated for 1 h on ice. Samples were sonicated with a Diagenode Biorupter on low setting for 30 s on and 30 s off for ten rounds at 4 °C and spun at max speed (21,130 g) for 10 min at 4 °C. 450 μl of sample were added to washed beads and incubated rotating at 4 °C for 2 h. Beads were washed 3x with IP buffer before proteins were eluted in 0.1 M glycine (pH 2.4) for 10 min at RT, and elution was quenched with an equal volume of 0.1 M Tris-HCl (pH 8.0).

#### Mass spectrometry sample prep

Immunoprecipitated samples were reduced with 10 mM DTT for 30 min at RT and alkylated with 20 mM iodoacetamide for 45 min at RT in the dark. Samples were adjusted to 10 mM CaCl_2_ and split into two aliquots. One set of aliquots was digested at RT with chymotrypsin at a ∼1:25 ratio and the other with trypsin at a ∼1:30 ratio. Digestions were quenched after ∼16 hrs by addition of TFA to pH 2. Samples were then desalted over Oligo R3 reverse-phase resin (Thermo Scientific, Cat#1-1339-03).

#### Data acquisition

Peptide quantification by LC-MS/MS was performed on a Thermo Fisher Ultimate 3000 Dionex^TM^ liquid chromatography system and a Thermo Q-Exactive HF-X^TM^ mass spectrometer. The mobile phases consisted of 0.1% formic acid aqueous (mobile phase A) and 0.1% formic acid 80% acetonitrile (mobile phase B) with a gradient of 5-45% over 48 min and a 60 min total gradient. Samples were quantified by A_280_ absorbance and 1 µg of each was injected. Trypsin samples were run with MS1 settings of 250-1100 m/z window, a resolution of 60,000, AGC target of 5e5, and MIT (maximum inject time) of 54 ms. MS2 scans were collected in data dependent mode with a TopN loop count of 10, resolution of 15,000, AGC target of 1e5, and MIT of 100ms. Chymotrypsin samples were run on the same LC gradient with MS1 settings of 250-1100 m/z window, a resolution of 60,000, AGC target of 1e6, and MIT of 60 ms. MS2 scans were collected in data dependent mode with a TopN loop count of 10, resolution of 15,000, AGC target of 5e5, and MIT of 120ms. Fragmentation was performed with HCD using stepped normalized collision energies (NCE) of 25, 27, 30%^95^.

#### Data processing

Data files were processed by Sequest^TM^ within Proteome Discoverer^TM^ (PD) 2.3 workflow nodes. Searching parameters were set to find mass offsets of 8.014 Da for heavy K(+8) lysine and 10.008 Da for heavy R(+10) arginine for the SILAC heavy pairs^55^. Additionally, phosphorylation (79.966Da) and methylation (14.015Da) were searched on both viral and host proteins. A human protein FASTA and adenovirus type 5 specific FASTA files downloaded directly from Uniprot were used to process the raw files^96^. No imputation was used across data files. A 1% FDR level cutoff was applied at the peptide level by Percolator and the protein level. The use of Minora Feature Detector^TM^ was used to identify SILAC pairs and identify non-sequenced peptides between runs^97^. Post-processing of the data files was performed in R Studio and peptide abundances were normalized to their respective proteins. If each peptide was identified in each sample, the heavy/light pairs p-values were determined by a Student’s t-test as previously used in both the original RBR-ID paper^53^ and the subsequent SILAC targeted RBR-ID paper^54^. Score plots and fold change plots were generated using the mapping function from the original RBR-ID paper^53^.

### hnRNP-C interactome analysis

In the targeted RBR-ID experiment, the “light” control sample contains the global interactome data for the targeted protein, hnRNP-C, in the absence of any RNA-protein crosslinking. The “light” control data for mock, Ad5 WT, and ΔE1B infections were compared to identify the proteins that interact with hnRNP-C in each of these conditions and to quantify changes in interactions induced by WT or ΔE1B infection. Three biological replicates were generated for each condition and each biological replicate was analyzed in two technical replicates. The “light” protein abundance data for each replicate was transformed by log2 and the identified protein abundances were normalized by the abundance of hnRNP-C in the respective replicate. The protein abundance values for the two technical replicates were averaged within each biological replicate. The protein abundance values for the three biological replicates within each condition were then averaged. When computing averages, unidentified values were not included in the calculation. Z-scores were calculated by comparing the average abundance of the protein in the respective condition to the mean and standard deviation of all averaged abundances for that condition. The z-score was calculated only proteins with abundance quantified in at least 2 replicates for the respective condition. Fold changes were calculated by comparing protein average abundance values for compared conditions. Hypothesis testing was performed by using an unpaired t-test to compare log2 normalized protein abundance values for each replicate within compared conditions. Correlations were analyzed for log2 normalized abundances using the cor function (Pearson correlation coefficient) and visualized using the corrplot package in the R software environment^98^.

### Statistics and reproducibility

Each experiment was carried out at least in triplicate with reproducible results. The sample size was chosen to provide enough statistical power apply parametric test (unpaired, two-tailed Student’s t-test unless otherwise noted). Details regarding statistical analysis are reported in each figure legend and p-values for each analysis can be found in **Supplementary Table 8**.

### Data availability

All mass spectrometry data for this study are deposited in the CHORUS database (dataset identifier and DOI will be provided upon acceptance of manuscript). The seCLIP-Seq data have been deposited in NCBI’s Gene Expression Omnibus^99^ and are accessible through GEO Series accession number GSE145411 (https://www.ncbi.nlm.nih.gov/geo/query/acc.cgi?acc=GSE145411). Additional supporting data are available from the corresponding author upon request.

### Code availability

The proteomics data were analyzed using standard methods. The implementation of the analysis was performed using R software. The scripts are available from the corresponding author upon request or can be accessed via GitHub. https://github.com/JosephDybas/AdenovirusProteomics

## Acknowledgements

We thank members of the Weitzman Lab for insightful discussions and input. We thank P. Choi for help with seCLIP-Seq, and G. Yeo and E. Van Nostrand for their input regarding analysis of corresponding data. We are grateful to A. Berk, P. Branton, R. Greenberg, G. Ketner, A. Levine, K. Lynch, D. Ornelles, J. Wilson, and H. Wodrich for generous gifts of reagents. We thank S. Schaffer for technical advice, and D. Avgousti, L. Busino, P. Choi, K. Lynch, and J. Weitzman for careful reading of the manuscript. We thank the UPenn Cell and Developmental Biology Microscopy Core for imaging assistance. This research was supported by NIAID grants R01-AI145266 (MDW), R01-AI121321 (MDW) and R01-AI118891 (BAG), and NCI grant R01-CA97093 (MDW). Additional support came from the NCI T32 Training Grant in Tumor Virology T32-CA115299 (AMP) and Individual National Research Service Awards F32-AI147587 (JMD) and F32-AI138432 (AMP) from the National Institutes of Health.

## Author Contributions

M.D.W., C.H., J.M.D. and J.C.L. conceived of the project. C.H. performed the experiments and received assistance from J.C.L., A.M.P. and E.T.K. C.H. prepared figures with input from other authors. J.M.D. performed the bioinformatics and proteomics data analysis. J.C.L. and R.L. performed mass spectrometry. A.M.P. performed RNA FISH. A.M.P., C.E.P., and C.H. performed RT-qPCR, CLIP-qPCR, and seCLIP-Seq sample preparation. K.E.H. performed bioinformatics analysis of seCLIP-Seq data. M.C. performed microscopy. C.H., J.C.L., and E.T.K. performed immunoprecipitation. C.H., J.M.D., and M.D.W. wrote the manuscript with input from the other authors. M.D.W. and B.A.G. supervised the research.

---

**Supplementary Figure 1.**
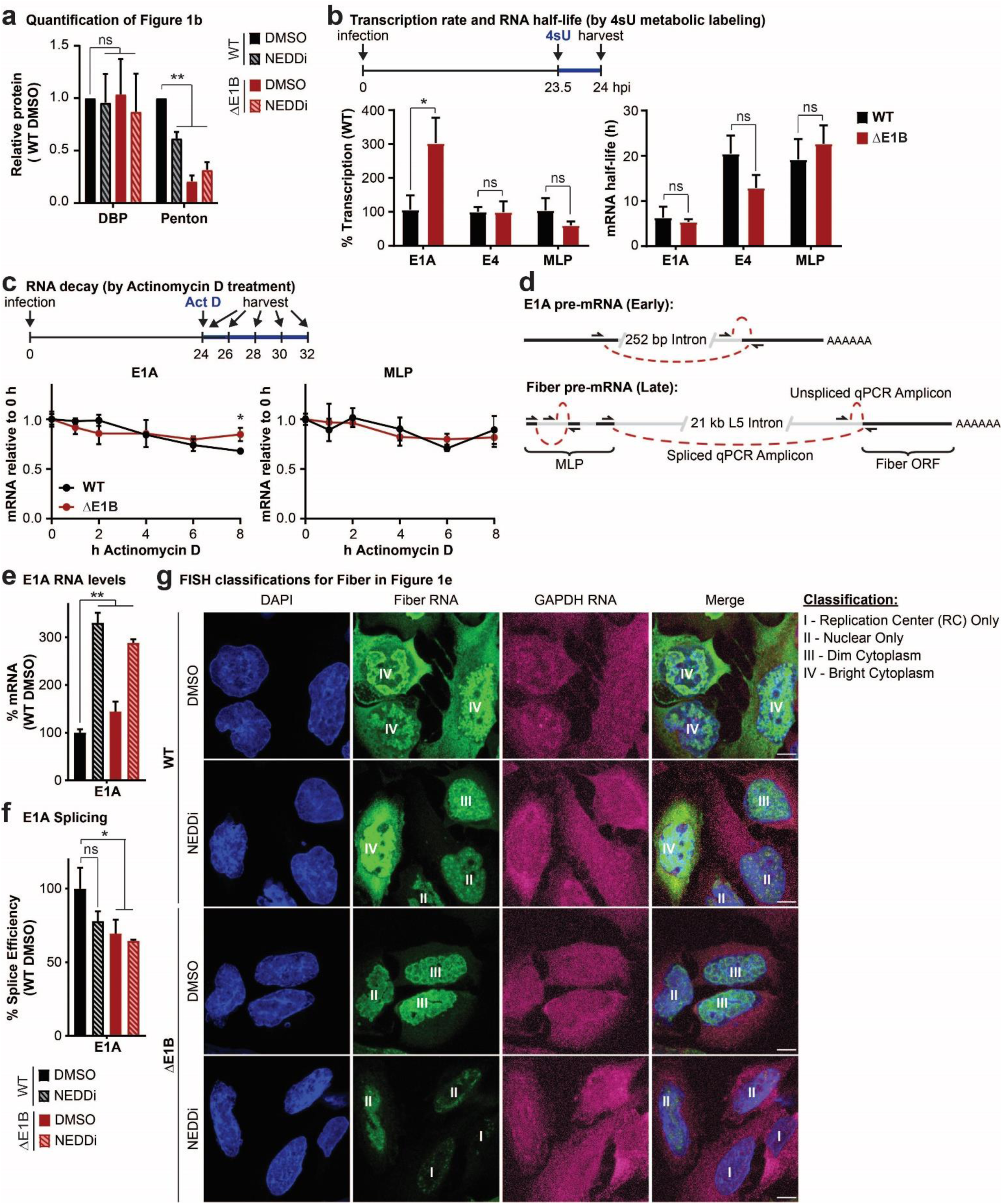
E1B55K deletion or inhibition of Cullin-mediated ubiquitination does not decrease late RNA transcription and decay or early RNA processing. **a,** Quantification of immunoblot shown in Figure 1b in triplicate. **b,** Analysis of nascent transcription and mRNA half-life by labeling RNA with 4-thiouridine (4sU) for 30 min at 23.5 hpi in HeLa cells infected with WT or ΔE1B Ad5 at MOI=10. Nascent 4sU-labeled RNA was purified for RT-qPCR for determining relative transcription rates of two early (E1A and E4) and one late (MLP) viral RNA. mRNA half-life was approximated using the ratio of nascent and total input RNA levels normalized to GAPDH. **c,** Analysis of decay of viral early (E1A) and late (MLP) RNA species by Actinomycin D pulse at 24 hpi in HeLa cells infected with WT or ΔE1B Ad5 at MOI 10 by normalization to input levels. **d,** Schematic illustrating primer design to differentiate spliced and unspliced viral transcripts. **e-g,** HeLa cells infected with WT or ΔE1B Ad5 (MOI=10) in the presence of DMSO or NEDDi (neddylation inhibitor MLN2449) added at 8 hours post-infection (hpi). Cells were harvested for RNA analysis at 24 hpi. **e,** Bar graph representing spliced RNA levels of viral early transcripts E1A by RT-qPCR, **f,** Bar graph representing splicing efficiency as the ratio of spliced to unspliced transcripts of E1A relative to the WT DMSO control by RT-qPCR. **g,** RNA FISH visualizing the localization of fiber (green) and GAPDH (magenta) transcripts in relation to nuclear DNA stained with DAPI (blue). Nuclei are labeled with the classification of each cell according to the pattern of fiber used for Figure 1d. Scale bar 10 μm. Shown is mean+s.d., n equals at least three biological experiments. Statistical significance was calculated using a paired (a) or unpaired (others), two-tailed Student’s t-test, * p < 0.05, ** p < 0.01.

**Supplementary Figure 2.**
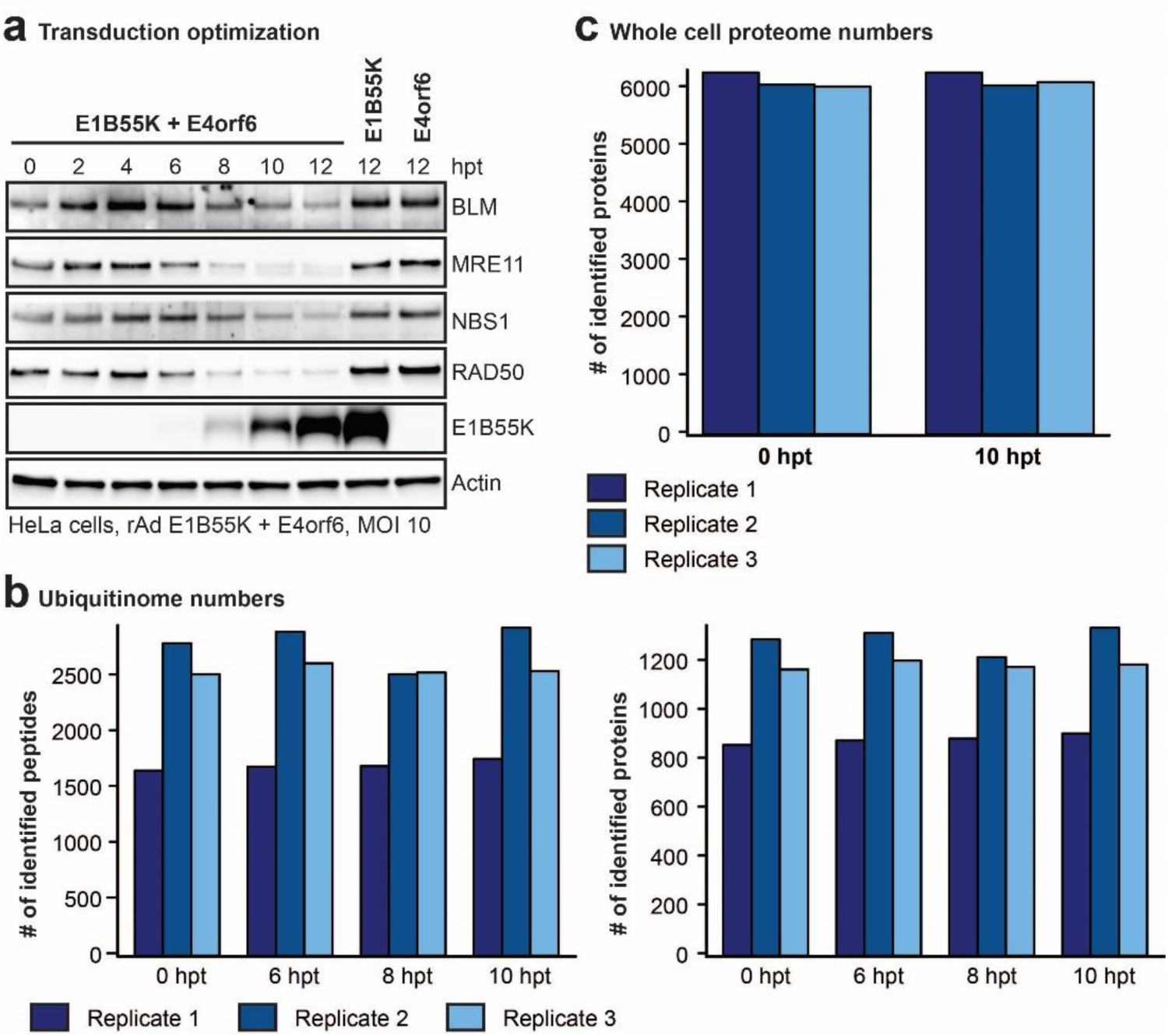
Quantification of number of peptides and proteins identified in di-glycine remnant profiling and whole cell proteome data sets. **a-c,** HeLa cells transduced with rAd E1B55K/E4orf6 at MOI 10. **a,** Immunoblot of time course of E1B55K/E4of6 expression showing degradation kinetics of known substrates. hpt = hours post transduction. **b,** Numbers of peptides and corresponding proteins identified following K-ε-GG antibody enrichment in di-glycine remnant combined with mass spectrometry analysis at 0, 6, 8, and 10 hours post E1B55K/E4orf6 expression. **c,** Number of proteins identified by whole cell proteomics analysis at time 0 and 10 hours post E1B55K/E4orf6 expression. **b,c,** Dark blue, medium blue, and light blue bars indicate the counts for three individual biological replicates.

**Supplementary Figure 3.**
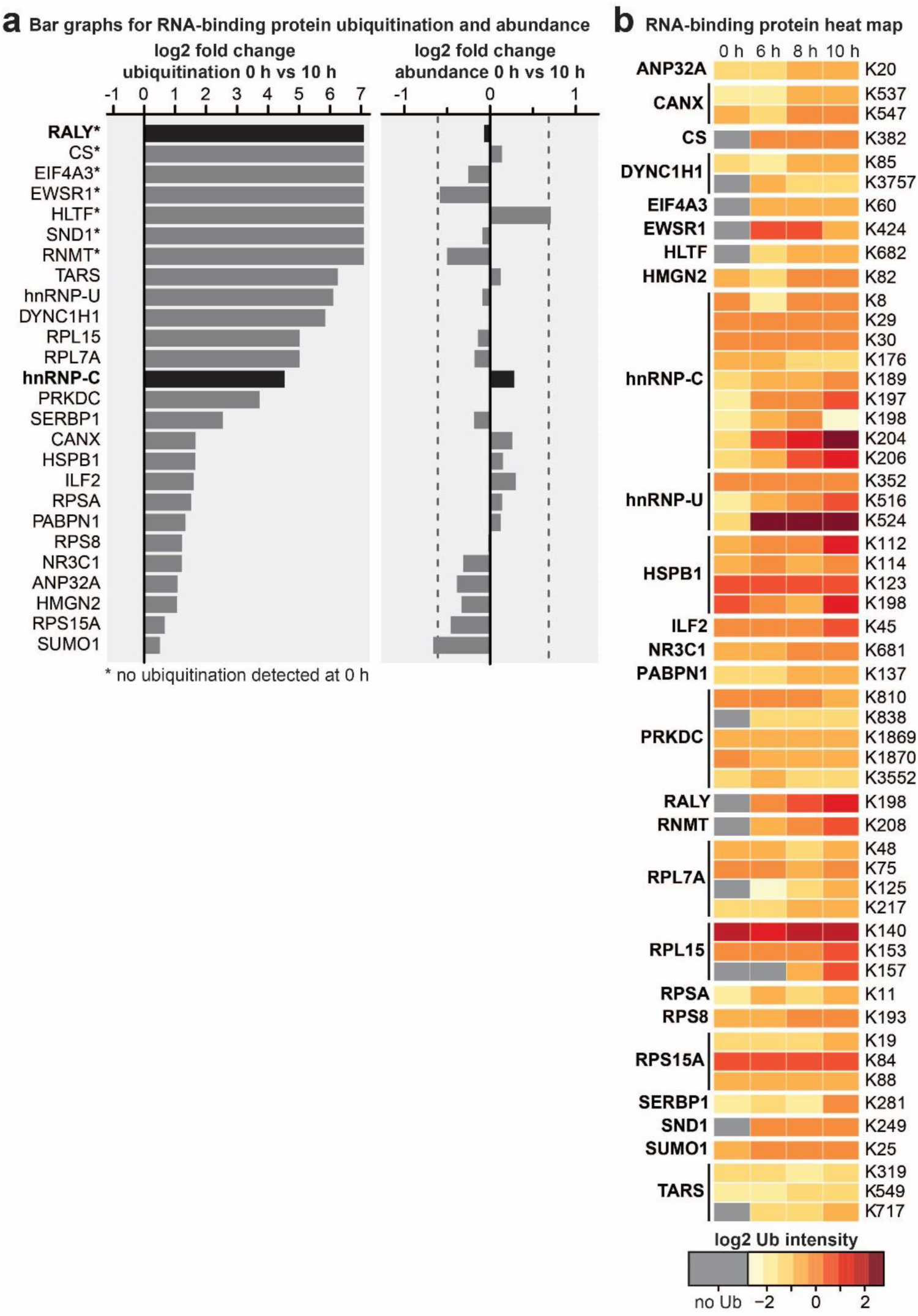
Di-glycine remnant profiling and whole cell proteome data for RNA-binding proteins enriched within the predicted E1B55K/E4orf6 substrates. **a-b,** Gene ontology analysis identified RNA-binding proteins enriched in the set of proteins that exhibited an increase in normalized protein-based ubiquitin abundance of log2 fold change > 1 following 10 h transduction of E1B55K/E4orf6. **a,** Enriched RNA-binding protein, ubiquitination log2 fold changes (left) and whole cell protein abundance log2 fold changes (right) following 10 h transduction by E1B55K/E4orf6. **b,** Heat map showing relative ubiquitination of the respective lysine residues quantified by di-glycine remnant profiling analysis at 0, 6, 8, and 10 h of E1B55K/E4orf6 expression for peptides within enriched RNA-binding proteins. Heat map color gradient is based on low (yellow) to high (red) ubiquitin abundance and grey indicates “not identified” at that time point.

**Supplementary Figure 4.**
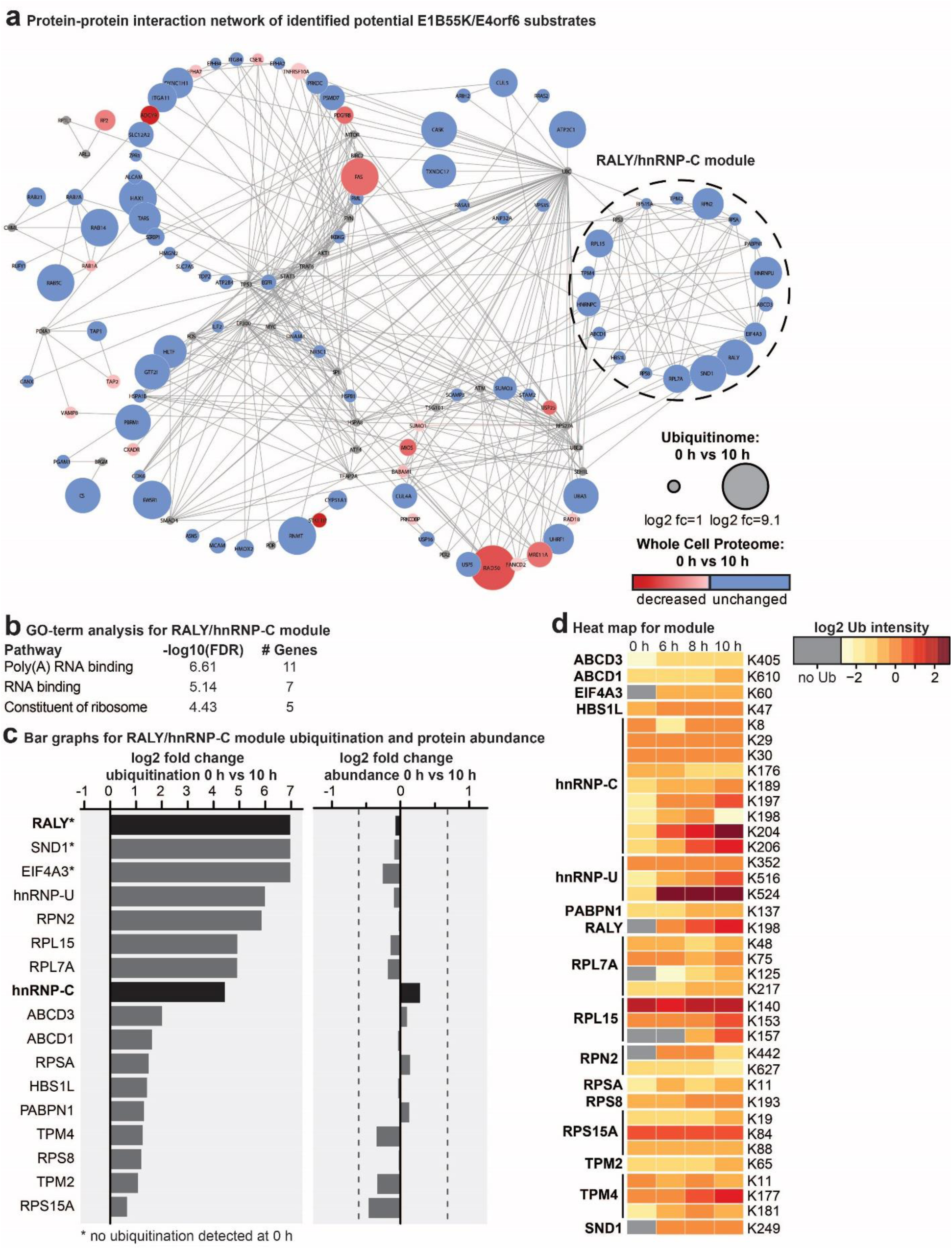
Network analysis of predicted E1B55K/E4orf6 substrates identifies a “RALY/hnRNP-C module” enriched for RNA-binding proteins. **a,** The Reactome-FI application in Cytoscape was utilized to generate a protein-protein interaction network in which nodes represent proteins and edges represent Reactome-based protein-protein interactions. Node size corresponds to relative protein-based ubiquitination log2 fold change and node color corresponds to whole cell proteome log2 fold change following 10 h transduction of E1B55K/E4orf6. Protein-protein interaction network of proteins that exhibited normalized protein-based ubiquitin abundance log2 fold change > 1 following 10 h transduction of E1B55K/E4orf6. Reactome-FI interaction module analysis was performed to generate clusters of highly interacting proteins. **b,** Gene ontology analysis for molecular function identified enrichment of RNA-binding and Poly(A) RNA-binding proteins within the RALY/hnRNP-C network module shown in Figure 2g. **c,** RALY/hnRNP-C network module protein ubiquitin log2 fold changes (left) and whole cell protein abundance log2 fold changes (right) comparing 0 and 10 h post-transduction with E1B55K/E4orf6. **d,** Heat map showing relative ubiquitin abundance quantified by di-glycine remnant profiling analysis at 0, 6, 8, and 10 h post E1B55K/E4orf6 transduction for peptides from proteins within the RALY/hnRNP-C network module. Heat map color gradient is based on low (yellow) to high (red) ubiquitination and grey indicates “not identified” at that time point.

**Supplementary Figure 5.**
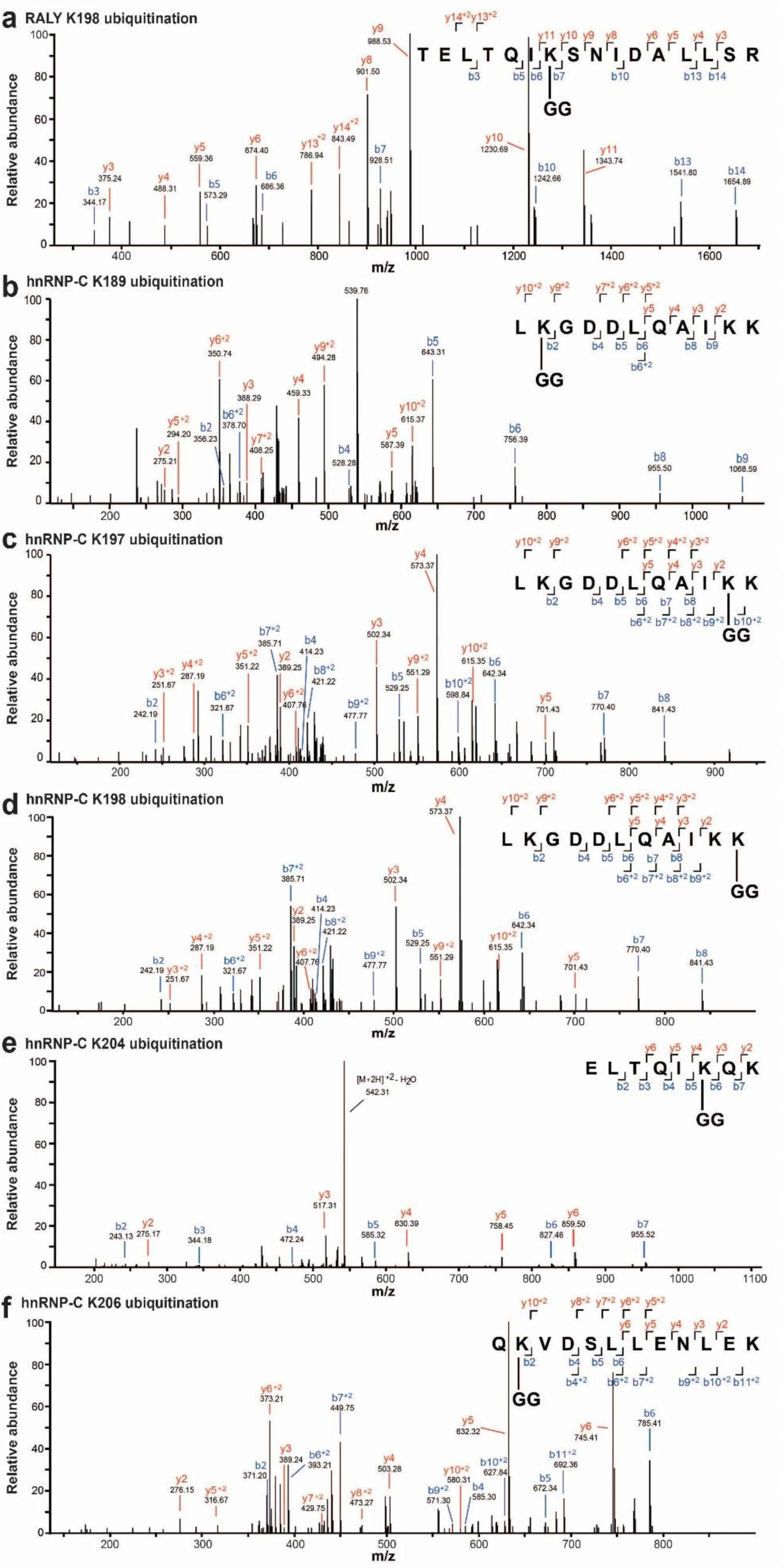
MS2 evidence for ubiquitination site localization in RALY (a) and hnRNP-C (b-f) peptides. Spectra were obtained from LC-MS/MS analyses using collision-induced dissociation (CID) at 35%, and identified in MaxQuant 1.6.0.1. All modified residues can be confidently identified by confirming ions, except for hnRNP-C K198 (d), which lacks ions to distinguish between K197 and K198. Best evidence spectra were selected for annotation of b-ion (blue) and y-ion (red) series and their masses for singly- and doubly-charged fragments.

**Supplementary Figure 6.**
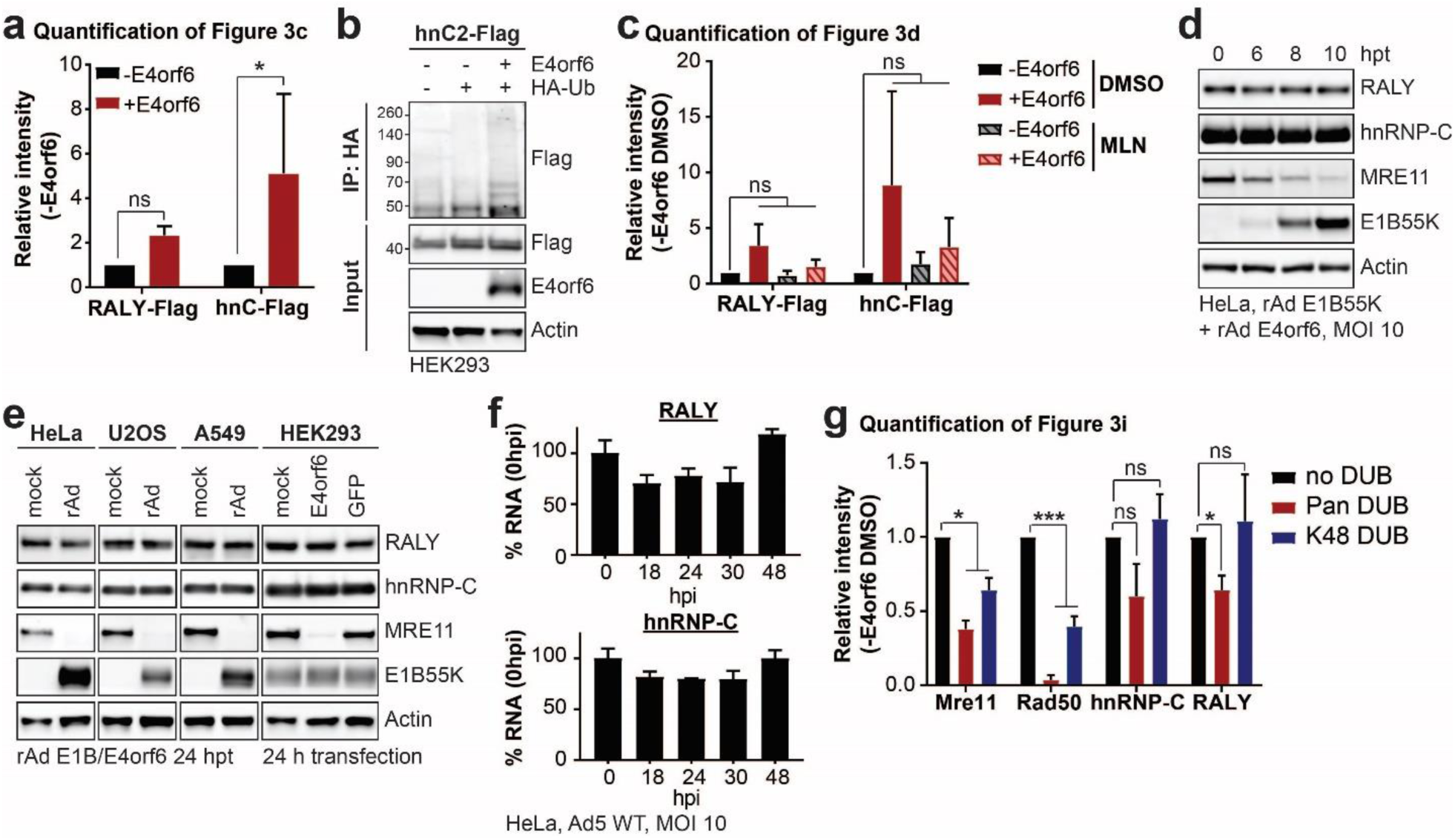
RALY and hnRNP-C are not decreased upon transduction in multiple cell lines. **a,** Quantification of immunoblot shown in Figure 3c in triplicate. **b,** HEK293 cells transfected with the indicated constructs for 24 h followed by denaturing IP with HA antibody and immunoblot analysis of hnRNP-C2-Flag. **c,** Quantification of immunoblot shown in Figure 3d in triplicate. **d,** Immunoblot analysis of protein levels in HeLa cells over a time course of transduction with recombinant Ad vectors expressing only E1B55K and E4orf6 (MOI=10). **e,** Immunoblot analysis of protein levels in HeLa, U2OS, A549 and HEK293 cells. HeLa, U2OS and A459 cells were transduced with recombinant Ad vectors expressing only E1B55K and E4orf6 for 24 h. HEK293 cells, which contain an endogenous copy of E1B55K, were mock transfected or transfected with plasmids expressing E4orf6 or GFP. **f,** Bar graphs of RALY and hnRNP-C RNA levels over a time course of infection with Ad5 WT (MOI=10) relative to mock as determined by RT-qPCR, shown is mean+s.d, n equals three biological replicates. **g,** Quantification of immunoblot shown in Figure 3i in triplicate. All immunoblots are representative of at least three biological replicates. Statistical significance was calculated using a paired, two-tailed Student’s t-test, * p < 0.05, ** p < 0.01, *** p < 0.005.

**Supplementary Figure 7.**
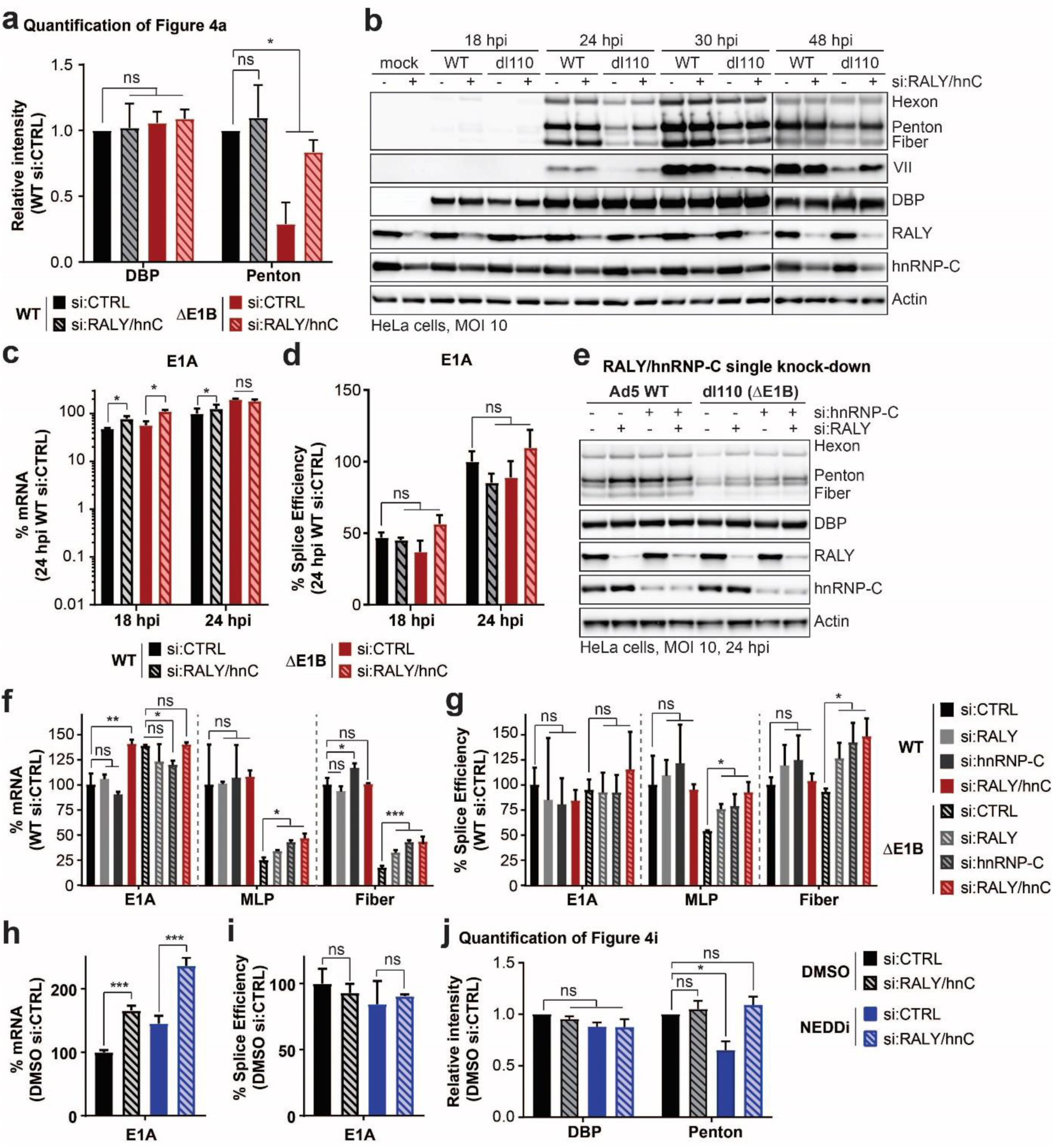
RALY and hnRNP-C single knockdown rescue late protein, RNA and splice efficiency during infection with Ad ΔE1B. **a-d,** HeLa cells transfected with control (siCTRL) or RALY and hnRNP-C (siRALY/hnC) siRNA 24 h prior to infection with Ad5 WT or ΔE1B (MOI=10), harvested at respective time points. **a,** Quantification of immunoblot shown in Figure 4a in triplicate. **b,** Extended immunoblot analysis of viral and cellular protein levels. **c,** Bar graph representing spliced RNA levels of viral early transcript E1A measured by RT-qPCR. **d,** Bar graph representing splicing efficiency as defined as the ratio of spliced to unspliced transcripts of E1A measured by RT-qPCR. **e-g,** HeLa cells transfected with control siRNA (siCTRL), siRNA for RALY (siRALY), siRNA for hnRNP-C (sihnRNP-C) or siRNA for both RALY and hnRNP-C (siRALY/hnC) 24 h prior to infection with Ad5 WT or ΔE1B (MOI=10) and harvested at 24 hpi. **e,** Immunoblot analysis of viral and cellular protein levels. **f,** Bar graph representing spliced RNA levels of E1A, MLP and fiber measured by RT-qPCR. **g,** Bar graph representing splicing efficiency as defined as the ratio of spliced to unspliced transcripts of E1A, MLP and fiber measured by RT-qPCR. **h-j.** HeLa cells transfected with control (siCTRL) or RALY and hnRNP-C (siRALY/hnC) siRNA 24 h prior to infection with Ad5 WT (MOI=10), treated with either DMSO or NEDDi at 8 hpi and processed at 24 hpi. **h,** Bar graph representing spliced RNA levels of E1A measured by RT-qPCR. **i,** Bar graph representing splicing efficiency as defined as the ratio of spliced to unspliced transcripts of E1A measured by RT-qPCR. **j**, Quantification of immunoblot shown in Figure 4i in triplicate. All immunoblots are representative of at least three biological experiments. All graphs show the mean+s.d. with n equals three biological replicates. Statistical significance was calculated using a paired (a and j) or unpaired (others), two-tailed Student’s t-test, * p < 0.05, ** p < 0.01, *** p < 0.005.

**Supplementary Figure 8.**
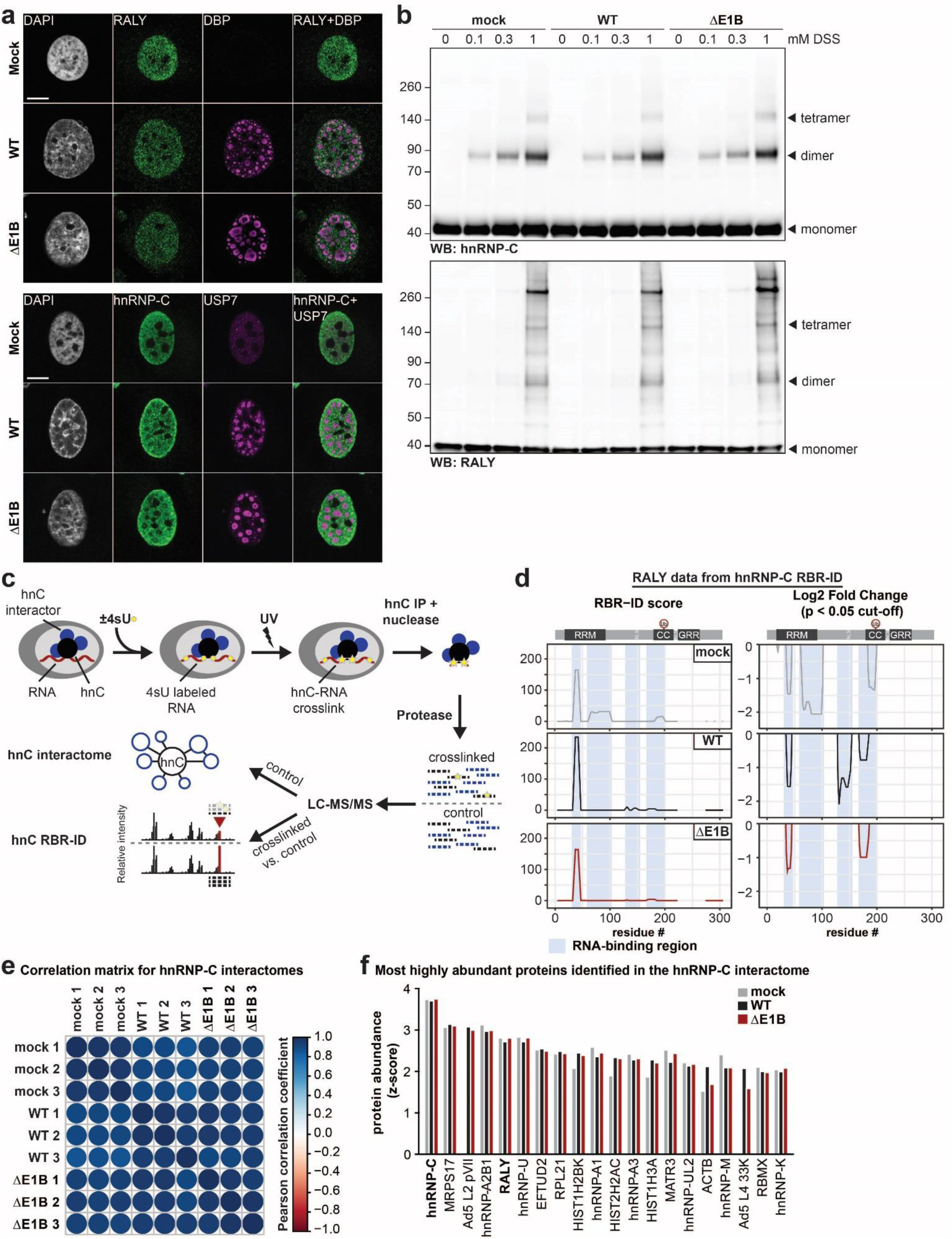
No dramatic difference in protein localization and protein-complex formation of RALY and hnRNP-C between Ad WT and ΔE1B infection. **a,** Representative images of immunofluorescence comparing the localization of RALY and hnRNP-C (both green) in mock, Ad WT and ΔE1B infection of HeLa cells (MOI=10, 24 hpi). Viral replication centers are stained by DBP or USP7 (both magenta) and nuclear DNA by DAPI (grey). Scale bar 10 μm. **b,** Immunoblot analysis of RALY and hnRNP-C protein complexes formed upon mock, Ad WT and ΔE1B infection of HeLa cells (MOI=10) and treatment with indicated concentrations of disuccinimidyl suberate (DSS) for 30 min at 24 hpi. Representative of three biological replicates. **c,** Schematic for targeted hnRNP-C RNA-binding region identification (RBR-ID) and interactome. **d,** Data for RALY from hnRNP-C RBR-ID experiment comparing mock (grey), Ad5 WT (black), and ΔE1B (red) at 24 hpi and MOI 10. Shown are smoothed residue-level RBR-ID score plotted along the primary sequence (**left**) and smoothed residue-level fold-change between crosslinked and control conditions with a significance threshold of p < 0.05 (**right**). RALY domain structure with ubiquitination site is shown above graphs. RNA-binding regions are highlighted in blue. **e,** Correlation matrix for hnRNP-C interactome between replicates of mock, Ad5 WT, and Ad5 ΔE1B. Color gradient is based on the Pearson correlation coefficient with correlation (> 0.0) in blue and anti-correlation (< 0.0) in red. **f,** Comparison of z-scores for top 20 proteins identified in hnRNP-C interactome during WT Ad5 infection (MOI 10, 24 hpi). Mock = grey, Ad5 WT = black, Ad5 ΔE1B = red.

**Supplementary Figure 9.**
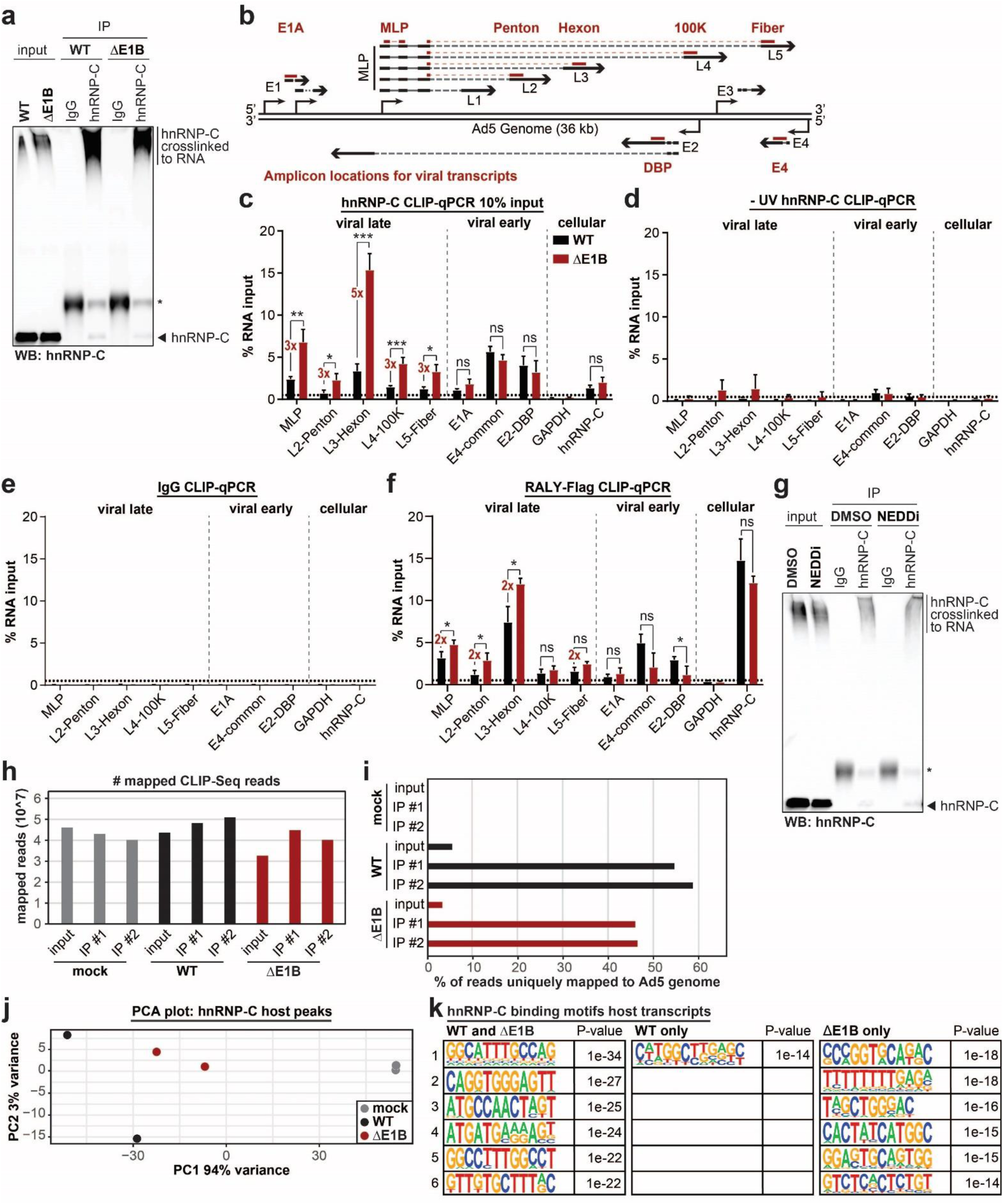
hnRNP-C and RALY interact more with viral late RNA in the absence of E1B55K. **a,** Control immunoblot for hnRNP-C CLIP-qPCR shown in Figure 5b. Higher molecular weight complexes stained with hnRNP-C antibody represent hnRNP-C crosslinked to RNA. * marks the antibody heavy chain detected in the IP. Representative of at least three biological replicates for both CLIP-qPCR and immunoblot analysis thereof. **b,** Schematic of the Ad5 genome and viral transcription units. Location of amplicons for viral early (E1A, DBP, E4) and viral late (MLP, Penton, Hexon, 100K, Fiber) are noted. **c,** HeLa cells infected with either WT Ad5 or ΔE1B (MOI=10), UV-crosslinked and harvested at 24 hpi, subjected to hnRNP-C CLIP with only 10% of input as compared to Figure 5b and RT-qPCR for viral early and late transcripts. **d,** HeLa cells infected with either WT Ad5 or ΔE1B (MOI=10), without UV-crosslinking and harvested at 24 hpi, subjected to hnRNP-C CLIP and RT-qPCR for viral early and late transcripts. **e,** HeLa cells infected with either WT Ad5 or ΔE1B (MOI=10), UV-crosslinked and harvested at 24 hpi, subjected to IgG CLIP and RT-qPCR for viral early and late transcripts. **f,** HeLa cells induced for RALY-Flag expression with doxycycline for 3 days total, infected with either WT Ad5 or ΔE1B (MOI=10), UV-crosslinked and harvested at 24 hpi, subjected to Flag CLIP and RT-qPCR for viral early and late transcripts. For all CLIP-qPCR experiments GAPDH is a cellular negative control. hnRNP-C is a cellular positive control. **g,** Control immunoblot for hnRNP-C CLIP-qPCR shown in Figure 5c. Higher molecular weight complexes stained with hnRNP-C antibody represent hnRNP-C crosslinked to RNA. * marks the antibody heavy chain detected in the IP. Representative of three biological replicates for both CLIP-qPCR and immunoblot analysis thereof. **h,** Number of mapped hnRNP-C eCLIP-Seq reads for the indicated conditions. **i,** Percentage of mapped reads that uniquely mapped to the Ad5 genome for the different hnRNP-C eCLIP-Seq conditions. **j,** PCA plot for hnRNP-C peaks mapped to host transcripts comparing mock (grey), Ad5 WT (black), and Ad5 ΔE1B (red). **k,** Top 6 hnRNP-C binding motifs identified for binding sites in WT and ΔE1B infection, WT infection only, and ΔE1B infection only.

## Notes

### Competing Interest Statement

The authors have declared no competing interest.

